# Emergent collective alignment gives competitive advantage to longer cells during range expansion

**DOI:** 10.1101/2024.01.26.577059

**Authors:** Nathan van den Berg, Kristian Thijssen, Thu Trang Nguyen, Mireia Cordero, Alba García Vázquez, Namiko Mitarai, Amin Doostmohammadi, Liselotte Jauffred

## Abstract

Bacterial competition shapes community architecture, yet a universally conserved determinant remains elusive. We show that cell aspect ratio –a simple morphological feature– confers a competitive advantage. Using growth-based range expansion experiments, we show that longer bacteria conquer the expanding front, even when initially in minority. Using an agent-based model of dividing bacteria, to isolate the effect of aspect ratio, we reveal that the takeover mechanism is collective alignment: groups of locally aligned bacteria form “nematic arms” bridging the central region of the colony to the expanding front. Once at the front, bacteria align parallel to it and block shorter bacteria from access to nutrients and space. We confirm this observation with single-cell experiments and further generalise our findings by introducing a generic continuum model of alignment-dominated competition, explaining both experimental and cell-based model observations. Moreover, we extend our predictions to spherical range expansions and confirm the competitive advantage, even though the effect is less pronounced than in surface-attached colonies. Our results uncover a simple, yet hitherto overlooked, mechanical mechanism determining the outcome of bacterial competition, which is potentially ubiquitous among various bacteria. Current advances in genetic engineering enable aspect ratio tuning as a mechanism with broad implications for biofilm control.

## INTRODUCTION

Bacteria’s competition for nutrients and territory drives biofilm evolution (1–4). The factors determining the outcome of competition among diverse bacterial species have a broad impact on a wide range of pathological (5), environmental (6), and microbiome interactions (7). While motility-related traits (8–11) and specific molecular mechanisms (12, 13) have been identified as potential winning attributes in bacteria.

Morphology is not always conserved from generation to generation, instead bacteria can change shape within a single generation, for example, during the infection of a host organism (14) or as a response to shifts in nutritional composition (15, 16), osmolarity (17), and temperature (18); allowing them to spread effectively. It has also been found that rod-shaped bacteria dominate the base of a colony, indicating that cell shape is an important determinant of biofilm structure (19). However, we still lack the full understanding of if and how cell shape affects competition during range expansion.

In this work, we find the aspect ratio to be a key determinant of competitive advantage. We show through range expansion experiments (20), how longer bacteria consistently dominate the colony front. Agent-based modelling reveals that this dominance arises from emergent collective alignment: radially aligned nematic sectors extend from the colony centre to the front, which they overtake by aligning and growing along it. These findings are supported by single-cell resolution experiments and a continuum model of alignment-driven competition. Furthermore, we did spherical range expansions (21) and concluded that the supremacy of longer bacteria is retained but weakened in a 3D configuration.

## RESULTS

### Long bacteria win exclusively – even when outnumbered

We began by mixing two substrains of the non-motile *Escherichia coli* B: wild-type (WT) with a plasmid coding for green fluorescent protein (GFP) and an *mreB* mutant (AK) with a smaller aspect ratio (Table 1 and Supplementary Fig. S1) and with a plasmid coding for red fluorescent protein (RFP), see Fig. 1A. The mixture was inoculated on agar (1.5%) and resulted (after incubation) in a colony with three distinct regions from the centre and outwards (Fig. 1B): (i) the “homeland” where both strains were well mixed, (ii) a band thought to arise from the up-concentration of cells on the rim of the inoculation droplet (i.e., “coffee ring” (22, 23) in Supplementary Fig. S2), followed by (iii) a band where sectors merged. Strikingly, the WT (green) strain won over the shorter AK (magenta) during range expansion (outermost region of iii). We verified that these strains had similar growth rates (Supplementary Fig. S3A) and that takeover was independent of inoculation density (Supplementary Fig. S4). However, we found that the shorter strain had a growth advantage (in respect to the longer) in co-cultures (Supplementary Fig. S3B-C). Thus, WT wins in surface-attached colonies, even though this strain has a fitness disadvantage in liquid cultures.

**Table 1:** Length (*l*), width (*w*), aspect ratio (*l*/*w*). *l* and *w* values are given as the mean ±SD (Supplementary Fig. S1).

| Name | $l$ (μm) | $w$ (μm) | $l/w$ |
| --- | --- | --- | --- |
| WT | $2.68 \pm 0.45$ | $0.68 \pm 0.09$ | 3.97 |
| AS | $2.34 \pm 0.34$ | $0.99 \pm 0.11$ | 2.36 |
| AK | $1.82 \pm 0.30$ | $1.05 \pm 0.13$ | 1.74 |

**Figure 1:**
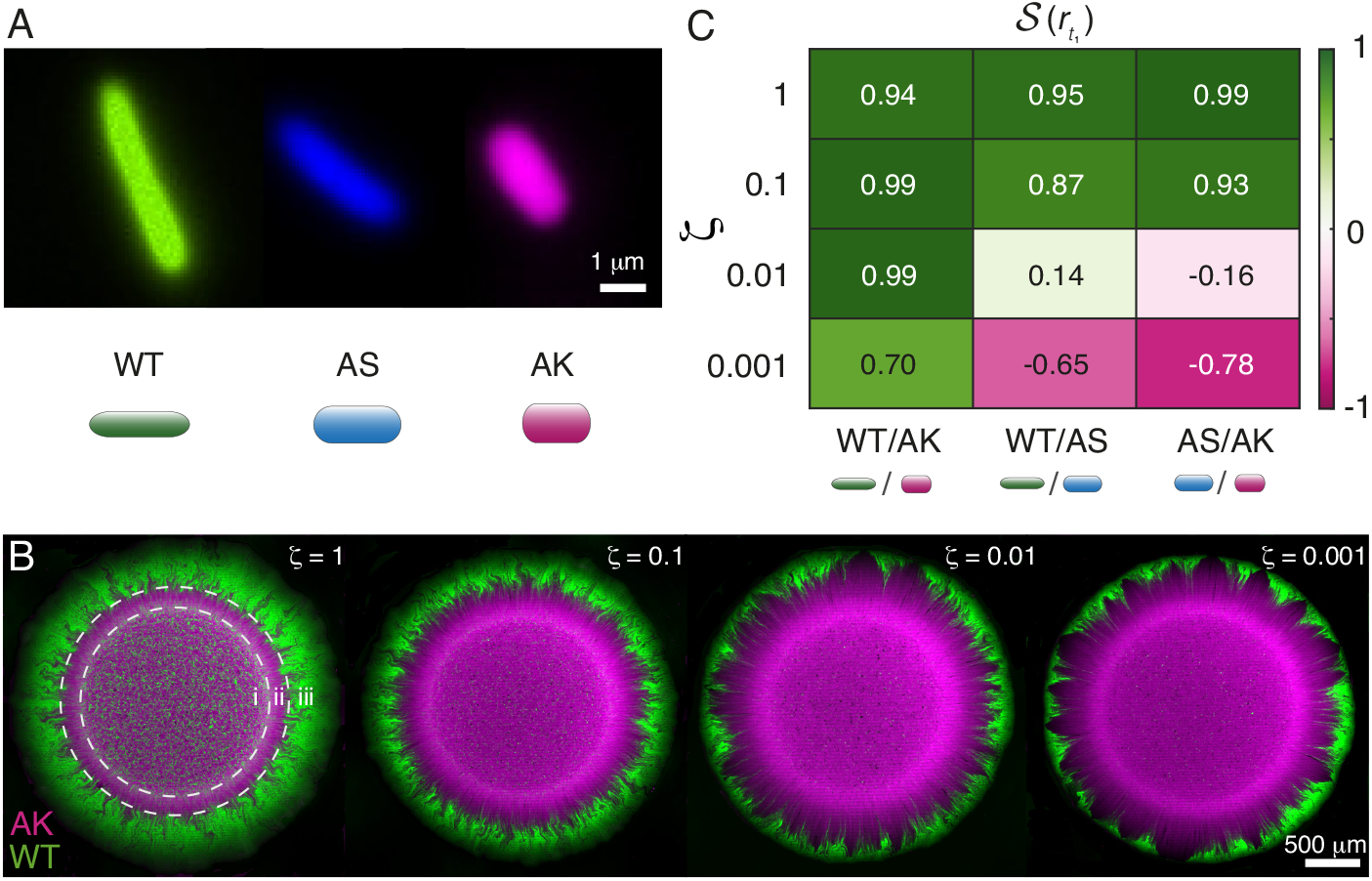
Long bacteria win even when initially outnumbered. **A:** Single-cell (pseudo-coloured) fluorescence images of the wild type (WT) and the *mreB* mutants (AS and AK). The scale bar is 1 µm. The images are representative images from Supplementary Fig. S1. **B:** Maximum-intensity projected (pseudo-coloured) confocal scanning laser microscopy images of 20 h competition experiments between WT (green) and AK (magenta) at different initial (WT/AK) ratios: ζ = {1, 0.1, 0.01, 0.001 }. Three distinct regions: (i) homeland, (ii) homeland rim, and (iii) a band where sectors merged. The scale bar is 500 µm. This experiment was repeated independently more than 3× for each ζ . **C:** Heat map of strength, 𝒮 (*r*_*t*1_), at the specific radial distance *r*_*t*1_ corresponding to the time: *t*_1_ = 1.5 days and averaged over several colonies (Supplementary Fig. S5). 𝒮 = − 1 corresponds to the longer cells (WT or AS) winning, 𝒮= −1 to the shorter (AS or AK), and 𝒮= 0 to equal strength (i.e., neutral fitness). The heat map shows all three strain combinations AS/AK, WT/AS, and WT/AK versus ζ . The cell graphics are to scale (Table 1).

To measure the strength of WT’s advantage, we assessed the competition outcome for different strain ratios ζ, which we defined as the inoculation density of the longer strain versus the shorter, quantified by optical density (OD_600_). We varied the ratio to identify the lower ζ-limit for WT to outcompete AK. However, for all tested strain ratios ζ (keeping combined inoculation density fixed), the longer strain (WT) consistently won over the shorter (AK) within the incubation time (20 h), see Fig. 1B. The takeover was noticeably slower in the ζ = 0.001 case, where WT sectors (green) spread azimuthally while curving around cone-shaped regions of shorter bacteria (magenta). Due to the difference in cell volume (Table 1), the number of cells in the stationary phase with same OD_600_ value is ∼ 1.3 times higher for WT than for AK (Materials and Methods). However, because WT outcompetes AK even for the low strain ratios (ζ ≪ 1), this slight advantage (in numbers) of WT is not enough to explain the competition outcome. Together, these results showed that the longer WT bacteria outcompete the shorter AK strain, even when initially outnumbered.

To explore the robustness of the impact of the aspect ratio, we introduced an additional *mreB* mutant (AS) with an intermediate aspect ratio (Fig. 1A and Supplementary Fig. S3 and set up competition experiments over 3 days (Supplementary Fig. S5. We kept the colour-coding, such that longer cells expressed GFP (WT or AS) and shorter RFP (AS or AK), and found no systematic dependence on the colour (Supplementary Fig. S6). While the intermediate AS strain lost against the longer WT bacteria, this previously losing AS outcompeted the shorter AK strain. This indicates that the aspect ratio of individual bacteria is the determining factor for the competition outcome in dividing *E. coli*. To quantify the competitive strength, we segmented and filtered the image stacks to obtain 2D masks reconstituting a connected colony surface, where all pixels were assigned one and only one colour. Behind the expanding colony front, additional cell-layers are formed, so we investigated the cross-sections (Supplementary Fig. S7) to rule out any dramatic changes in the composition of cells along the vertical direction. On this mask, we defined the radial distance *r* (*t*), from the homeland boundary (i.e., the ring of high intensity of shorter cells). Then, we measured the strength 𝒮 (*r*), defined by the re-scaled occupancy of the longer cell, such that 𝒮= 1 ( 𝒮= − 1) signified that the longer (shorter) cells fully dominated. Fig. 1C is a heat map of how ζ controls the strength, 𝒮(*r*_*t*1)_, at the radial distance corresponding to the time point *t*_1_ = 1.5 days of incubation (Materials and Methods). The quantitative analyses confirmed that WT always wins over AK, even when initially outnumbered by as much as ζ = 0.001 (i.e., one in thousands). For AS/AK and WT/AS combinations, we found few successful formations of sectors of the longer strain for ζ ≤ 0.01, indicating that the outcome is highly stochastic for small ζ (Supplementary Fig. S5); averaging over several colonies resulted in 𝒮(*r*_*t*1)_ ≲ 0. However, the sectors with longer bacteria tended to spread azimuthally with increasing *r*, indicating that they are likely to win the expanding front later on. In other words, the strain of longer aspect ratio could take over, if they succeeded in forming sectors early on (i.e., at small *r*).

### Alignment-induced mechanism gives longer bacteria a mechanical winning advantage

The experimental results suggested cell aspect ratio to be the determinant of the outcome of bacterial competition in the monolayer front. In order to test if this is solely based on the mechanics of bacterial organisation, we simulated the range expansion in a minimal agent-based model (Materials and Methods), where surface adhesion dominates. Bacteria were modelled as non-motile repulsive particles expanding on a 2D surface. The particles grew with the same fixed rate (with stochasticity). We distinguished longer from shorter species by setting the length at division as *l*_*L*_ and *l*_*S*_, respectively. By using a constant width of *w* = 1, the aspect ratio (*l*/*w*) of these particles were *l*_*L*_ and *l*_*S*_. We used an inoculation ratio, ζ_*i*_ = 0.25 (ratio of number of long species over short), such that shorter bacteria were in majority, and placed individual particles (either *l*_*L*_ = 8 or *l*_*S*_ = 3.5) randomly in a small region. We then let the cells grow and interact to expand over the 2D surface, as seen in Fig. 2A (Movie 1).

**Figure 2:**
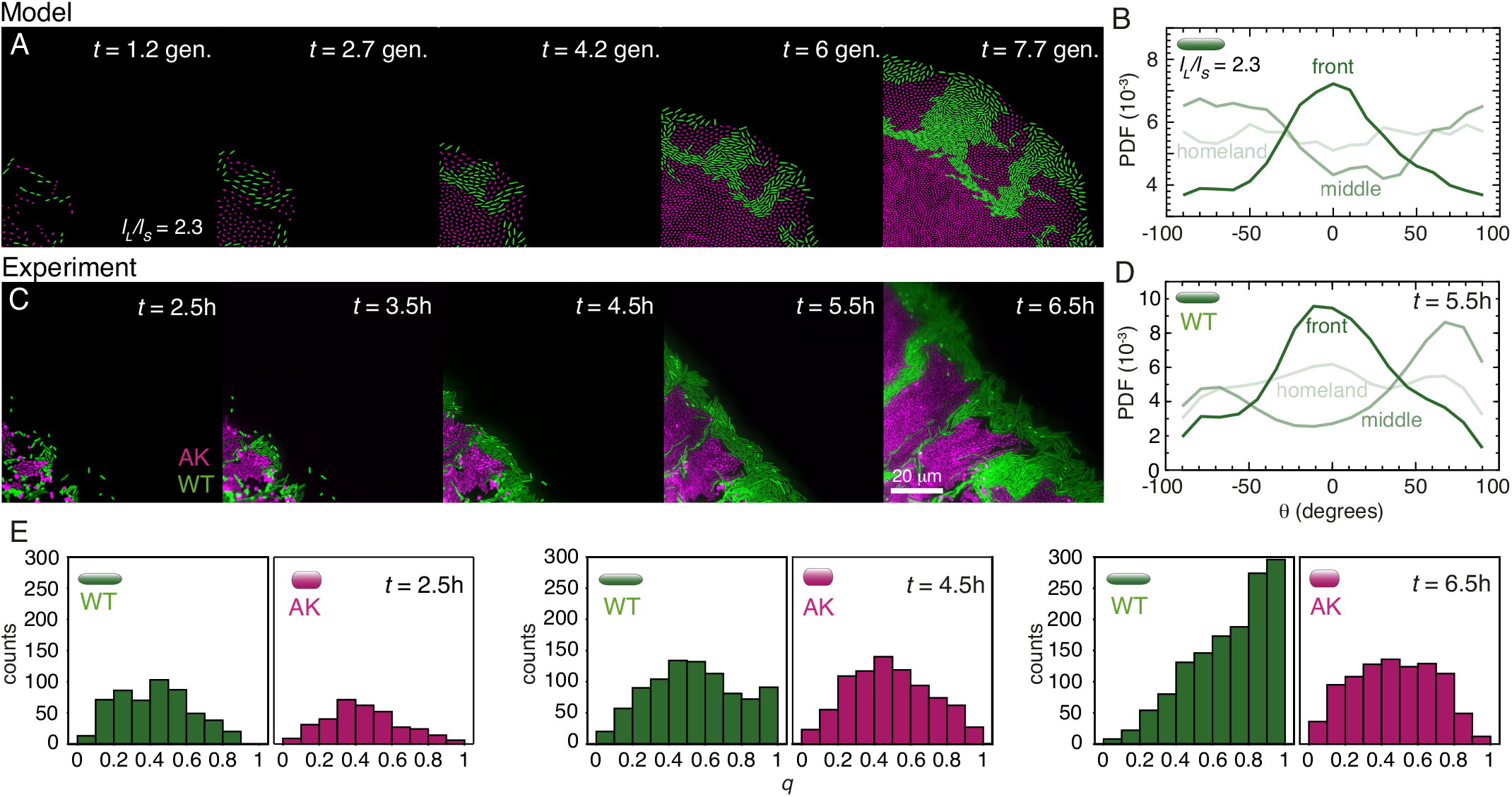
Alignment-induced mechanism gives longer bacteria a mechanical winning advantage. **A:** Time series of particle-based simulations, where long bacteria (green) overtake the short bacteria (magenta) at the colony front by forming expanding regions along the front. The ratios between the aspect ratios were *l*_*L*_/*l*_*S*_ = 2.3 ( ∼ aspect ratio difference of WT and AK) and between the species: ζ_*i*_ = 0.25. The full time-lapse is given in n 1. **B:** The probability distribution function, PDF, of the angle, *θ*, to the expansion front at three equidistant regions of *r*: the region were particles originally was deposited (homeland), closest to the outer rim (front), and the region in between (middle) at time: *t* = 7.7 gen. **C:** Snapshots from (pseudo-coloured) fluorescence time-lapses of competition between WT (green) and the *mreB* mutant AK (magenta) at time *t* after inoculation. The scale bar is 20 µm. The full time-lapse is given in Movie 2. **D:** The probability distribution function, PDF, of the angle, *θ*, of WT to the expansion front at three equidistant regions of *r*: where the inoculation droplet originally was deposited (homeland, *N* = 351), closest to the outer rim (front, *N* = 385), and the region in between (middle, *N* = 338). Data are from *t* = 5.5 hours in (C), like the AK PDF (Supplementary Fig. S8). **E:** The nematic order parameter, *q*, of the experimental data in (C) for WT and AK individually corresponding to every second time frame in (A) *t* ∈ {2.5, 4.5, 6.5 h. The distributions did not differ significantly after 2.5 h (*p* > 0.1) but they differed increasingly after 4.5 and 6.5} h (*p* < 0.001).

After the homeland had reached uniform density and the radial expansion began, sector boundaries between populations are formed. Active forces cause elongated bacteria to align along these sector boundaries, a process known as active anchoring (24–27). These interface boundaries orient the elongated bacteria into narrow channels (28) along the radial vector, **r**, which connect the interior (homeland) to the outer rim of the colony, forming conduits through which the longer bacteria can reach the front. This channel-formation resulted in azimuthal spread of sectors of longer bacteria with increasing radial distance from the centre of the inoculation to the expanding front, *r* _*f*_ . As a result, the longer bacteria are deposited at the front, where they form localized bumps (*t* > 4 gen. in Fig. 2C and Movie 1). Once in front, they spread out (29) and active forces result in a predominant azimuthal alignment (along the front) of the longer bacteria, as observed in Fig. 2B. See Supplementary Information for details on active anchoring. This alignment forms a barrier that suppresses the further expansion of the shorter bacteria. For comparison, we simulate a case in which short cells divide faster (Supplementary Fig. S9), and observe that this barrier does not form in that scenario.

To test the mechanism suggested by the cell-based model and investigate how populations compete at the individual cell scale, we ran time-lapse experiments with single-cell resolution. *E. coli* colonies were inoculated from a suspension of WT/AK in equal proportions (ζ = 1). We imaged the growth at the colony front over time *t*, as shown in Fig. 2C (Movie 2) and for additional time-lapses (Supplementary Fig. S10-11, Movie 3-4). Individual bacteria were segmented and their orientation was determined and the angle, *θ*, to the expanding colony front (i.e., 90^°^ to **r**) in three equidistant regions of *r* was measured (Fig. 2D). In agreement with the alignment-induced mechanical mechanism suggested by the agent-based model, we found that the winning WT bacteria align along **r** (inside the colony) or along the expanding colony front, once the cell monolayer becomes confluent (Supplementary Fig. S11). Furthermore, the nematic order parameter, *q*, which is a measurement of local orientation alignment (30), was calculated for every bacterium relative to its 8 closest neighbours, as extensively reported for bacterial mono-cultures (8, 31–35). The distribution of *q* was analysed at three different time points: *t* ∈ {2.5, 4.5, 6.5} h in Fig. 2E. The distributions of WT and AK strains did not differ significantly after 2.5 h (*p* > 0.1), but they differed increasingly after 4.5 and 6.5 h (*p* < 0.001). The distributions account for the visual observations in Fig. 2C that WT cells align with each other, while AK’s alignment was weak. Together, model predictions and experimental quantifications show how the longer WT are squeezed by small clusters of growing AK cells and form channels of (radially) aligned bacteria. When they reach the colony front, they align with the expanding front and start dividing along the tangential direction. In this way, WT cut their shorter AK counterparts off from both nutrients and space.

To emphasise the generic mechanical nature of this takeover mechanism – set by aspect ratio affecting torques and enhanced by stronger local alignment of the longer bacteria – we also employed a continuum model of a bi-phasic active nematic, where we took the two different bacteria types into account with an additional phase field order parameter *𝜙*, where *𝜙* > 0 (*𝜙* < 0) corresponded to longer (shorter) bacteria regions modelled with a higher (lower) elastic constant and extensile activity (which mimics the dipolar force of division, without changing the actual biomass at the front (36, 37)). It is well established that both orientational elasticity and extensile division activity are higher for higher aspect ratios (38, 39). We observed that even when the front is mostly populated by shorter bacteria, the phase corresponding to the more elongated bacteria eventually overtakes the shorter (Supplementary Fig. S12). Importantly, this phase segregation is solely driven by activity, stemming from cell divisions, and differences between aspect ratios, without any free-energy-driven phase separation. Remarkably, the bacteria of longer phase take over the front by forming aligned channels to the interfaces, due to differences in torque and local alignment resulting in “active anchoring” at the front (24), in agreement with both the cell-based model and experimental observations (Fig. 2A and 2C). Consequently, the bacteria of longer phase spread at the interface, as they aligned more strongly with it, due to their higher elastic constant. We note that no radial expansion occurs in this model, and thus no radial sector boundaries are present for radial alignment. Instead, bacteria with longer phase exhibit increased activity due to reduced friction, leading to active mixing in which the longer bacteria are driven toward the interface (40). As the total biomass is preserved in this minimal model, we highlight a distinct collective mechanical mechanism, which induces the taking over of the longer bacterial phase, regardless of any possible number advantage in inoculation.

### Takeover rate is set by differences in aspect ratios and the length of the shorter cell

Having established the competitive advantage of a large aspect ratio through experimental observations, cell-based and continuum models, we next turned to investigating the takeover rates. Fig. 3A shows the experimentally obtained strength 𝒮 (*r*) from the homeland (*r* = 0) to the outer rim of the colonies for the strain combinations and ζ -values that clearly switched within 3 days (i.e., 𝒮 (*r*_*t*1)_ > 0.85 in Fig. 1C). We defined the takeover rates, *v*, as the slope of 𝒮 (*r*) and summarised them in Fig. 3B versus the length of the shorter strain (i.e., *l*_*S*_). We also measured the development of 𝒮 over time by following the front and confirmed that the takeover rate *v* is consistent with the dynamics (Supplementary Fig. S13). We paralleled this result with the agent-based simulations to measure the time-dependent strength, 𝒮 (*r* _*f*_ (*t*)) (Fig. 3C) and the associated takeover rate, *v*_sim._. This rate was also dependent on the aspect ratio difference *l*_*L*_/*l*_*S*_, as a more significant *l*_*L*_/*l*_*S*_ resulted in a stronger alignment along the channels (Supplementary Fig. S14). Hence, the longer bacteria had enhanced access to the colony front. This is in agreement with the experimental observation that *v* is larger for WT/AK than WT/AS.

**Figure 3:**
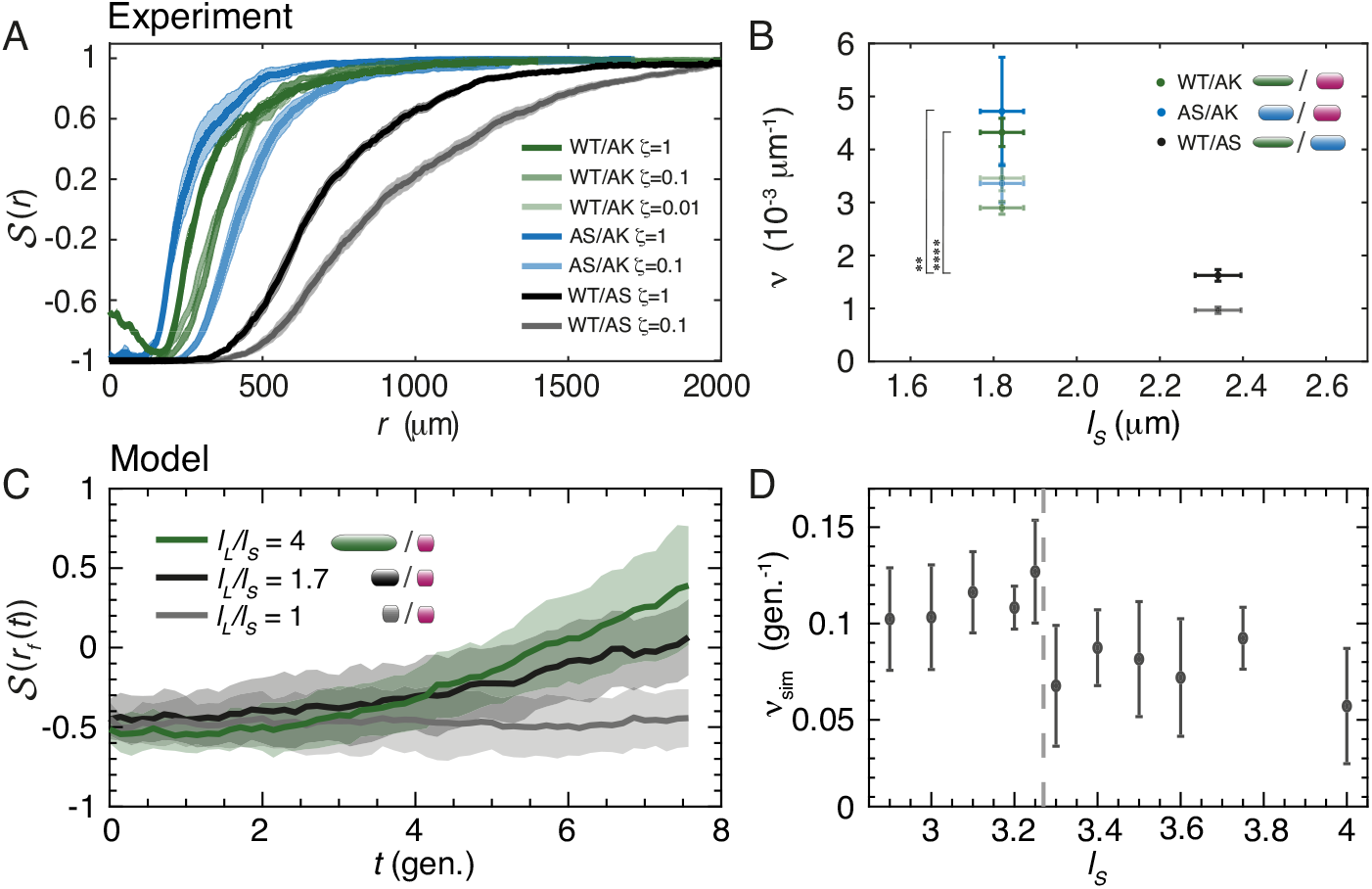
Takeover rate is set by differences in aspect ratios and the length of the shorter cell. **A:** Competition is evaluated by the strength, 𝒮 (*r*), along the radial length, *r*, for the strain and ratios, ζ, and strain combinations WT/AK (green), AS/AK (blue), and WT/AS (grey). We collected data from *N* ∈ {5, 4, 5, 6, 8, 9, 9} colonies (following order of legend) and ran the experiments for 3 days for all colonies to switch completely from − 1 to +1, which was the case for strain combinations and ζ -values that fulfilled 𝒮 (*r*_*t*1)_ > 0.85 in Fig. 1C. Notice how WT/AK mixed ζ = 1 drops from a higher 𝒮-level at the homeland boundary (*r* = 0) before switching again. The shaded regions correspond to ±SEM. **B:** Takeover rate, *v*, from for WT/AK (green), AS/AK (blue), and WT/AS (grey) versus length of the shorter cell, *l*_*S*_, for varying ζ (opacity as legend in (A)). Slopes are found for the individual colonies by fitting the linear part around 𝒮 ∼0 and *v* is the ensemble-average over *N* ∈ {5, 4, 5, 6, 8, 9, 9} colonies (following order of legend in (A)). The error bars correspond to ±SEM. *v* of WT/AS (ζ = 1) is significantly different from AS/AK (^∗∗^, *p* = 0.0025, two-sided T-test) and WT/AK (^∗∗∗∗^, *p* = 1.2 · 10^−7^, two-sided T-test). **C:** The time evolution of strength at the front, 𝒮 (*r* _*f*_ (*t*)), where *r* _*f*_ (*t*) is the radial distance from the centre of the inoculation to the expanding front at time, *t*. The long bacteria cell aspect ratio is fixed to *l*/*w* = 4, the species ratio is set to ζ_*i*_ = 0.25, and *l*_*L*_/*l*_*S*_ = 4, 1.7, and 1 are shown. The shaded regions correspond to ±SEM and *N* = 10. **D:** The simulation takeover rate, *v*_sim._, (slope of (C) between *t* = 4 gen. to *t* = 8 gen.) for varying lengths of the shorter bacteria, *l*_*S*_, with fixed ratio between long and short bacteria: *l*_*L*_/*l*_*S*_ = 1.5. In simulations, bacteria with *l*_*S*_ ≤ 3.25 (dashed vertical line) are dominantly isotropic and bacteria with larger *l*_*S*_ are dominantly nematic. The error bars correspond to ±SEM and *N* = 10.

Surprisingly, we found that the takeover rates for the competition between longest and intermediate strains, WT/AS (grey), are significantly slower than for the two other strain combinations (longest versus shortest, WT/AK, and intermediate versus shortest, AS/AK), although the ratios of aspect ratios (Table 1) were similar for WT/AS and AS/AK (∼ 1.5). Since our results established that the alignment-induced mechanism governs the competition between growing bacteria, we hypothesised that the difference in takeover rates were related to the relative ability of bacteria to form nematic domains. To explain the differences in the takeover rate *v* between WT/AS (longest versus intermediate) and AS/AK (intermediate versus shortest) (Fig. 3B), we used the agent-based model to examine the takeover rate for fixed *l*_*L*_/*l*_*S*_ ratio ( ∼ 1.5), while varying *l*_*S*_ from no local alignment (isotropic) to local alignment (nematic). The transition between isotropic and nematic states is set by excluded volume interactions and is dictated by local density and the particle aspect ratio, with corrections predominantly due to polydispersity of the length. As a result, we absorb the width variations of the bacteria strains in the particle aspect ratio (41, 42). Given our simulation doubling time, the isotropic-nematic transition occurred around length *l*_*S*_ = 3.25. The agent-based model showed that the takeover rate dropped when both bacteria strains had local orientation ordering, see Fig. 3D. This explains why the switching speed was slower for WT/AS than for AS/AK: WT and AS both had nematic order, while in the AS/AK strain combination, only the longer AS were nematically aligning. This is also in accordance with the observation that the sector boundaries are more rugged for WT/AS (Supplementary Fig. S5), indicating that AS cells might be weakly aligning. Together, these results further highlight the directed radial alignment (leading to takeover) results from the interplay between nematic and non-nematic bacteria. Specifically, nematic bacteria align at the boundaries formed with isotropic bacteria and this alignment must be diminished, when neighbouring bacteria are weakly nematic themselves. The reason is that active anchoring becomes weaker, as it scales with the nematic gradient at the interface (24). For instance, the radial expansion of isotropic AK patches directs the chain-formation of WT into the nematic arms (Fig. S10). Furthermore, the takeover rate is independent of the length of the shorter bacteria, in this case of an isotropic and nematic-packing species (Fig. 3D). However, if the shortest bacteria species transition from isotropically packed to nematically ordered state, the takeover rate drops. This radial alignment effect disappears in single species colonies (Supplementary Fig. S15 and Fig. S16), as the incompressible radial expansion flow, in the absence of interspecies interfaces, does not induce radial alignment (43, 44).

### Competition strength weakens in 3D

After establishing the significant advantage of elongation for surface-attached cells, we turned to 3D growth. The results so far have established that emergent nematic alignment is a competitive advantage. The local nematic order has been observed in 3D biofilms (45), though theoretically, it is expected that the nematic alignment weakens in 3D (46, 47) and as such we expected that the takeover strength weakens, compared to surface-attached (2D) conditions. We tested our predictions in WT/AK colonies grown from inoculation beads of small volume (0.05 nl) in low-density (0.5%) agar (inset of Supplementary Fig. S17). We defined a strain ratio, ζ_*n*_, as the ratio of colony-forming units (CFU) of the longer strain versus the shorter (Supplementary Fig. S17), and imaged (with confocal-scanning laser microscopy) the resulting 3D colonies for various ζ_*n*_, as shown in Fig. 4A. We then sorted the ensemble of colonies into subgroups of either mono-coloured colonies (WT or AK) or colonies with both strains surviving at the surface (grey); distributions are given in Fig. 4B. For comparison, we also calculated (assuming a binomial distribution) the fraction of colonies that initially contained only AK or WT. The probability of a pure WT inoculation was negligibly small in all cases, in contrast to the probability of pure AK. It is, thus, unlikely that all mono-coloured WT colonies arose from pure WT inoculation beads. Instead, the longer strain (WT) had outcompeted the shorter (AK) during range expansion. We also found examples of the inverse, where AK had taken over the colony surface, even though the inoculum had both WT and AK.

**Figure 4:**
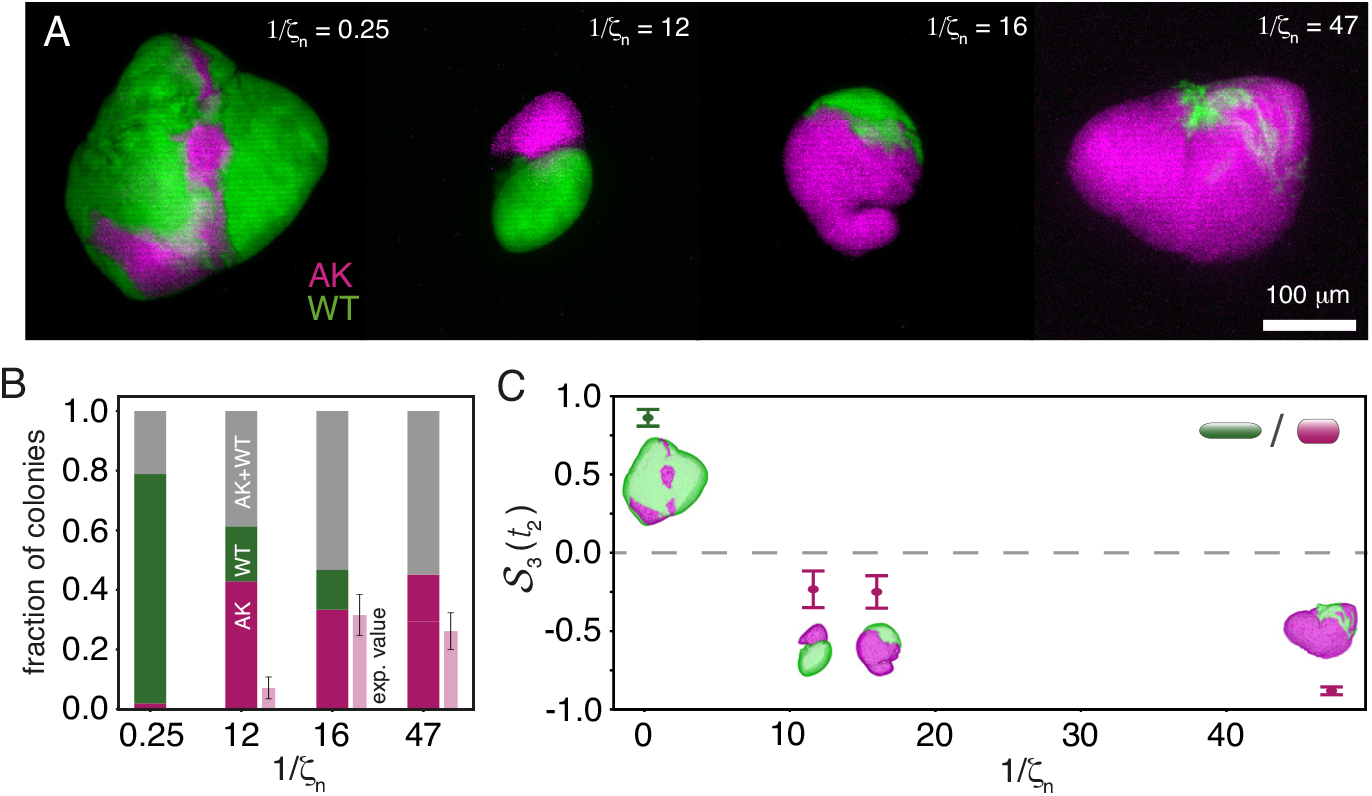
Competition strength weakens in 3D colonies compared to surface-attached bacteria. **A:** Maximum-intensity projected (pseudo-coloured) fluorescence images of colonies formed from WT/AK inoculation beads of different ratios, ζ_*n*_. The scale bar corresponds to 100 µm. **B:** Fraction of colonies with mono-coloured surfaces (WT or AK) or both (WT+AK). Data are presented as fractions ±SD and the number of colonies were *N* ∈ {52, 49, 45, 53} for 1/ζ_*n*_ ∈ {0.25, 12, 16, 47}, respectively. Predicted number of colonies (exp. value) originating from pure AK inoculation beads, using the numbers from Supplementary Fig. S17. Data are presented as fractions and error bars correspond to ±SD. **C:** Strength, S_3_(*t*_2_), of the ensemble of colonies versus 1/ζ_*n*_, measured in (A). Here 1 (−1) corresponds to the longer (shorter) cells winning within the time, *t*_2_ = 10 h, and 0 to equal strength (dashed grey line). The error bars correspond to ±SEM and the number of colonies were *N* ∈ {52, 49, 45, 53} for 1/ζ_*n*_ ∈ {0.25, 12, 16, 47}, respectively.

To quantify the takeover strength, we segmented and filtered the image stacks to obtain 3D masks reconstituting a connected colony surface with the width of a single voxel, which were assigned one and only one colour (Supplementary Fig. S18 and Movie 5). In order to determine the strength, 𝒮_3_ (*t*_2)_, at a specific time, *t*_2_, we quantified the proportion of the WT strain on the colony surface, as found in Fig. 4C. Interestingly, we found that when WT/AK are mixed four to one (i.e., 1 / ζ_*n*_ = 0.25) WT occupies > 90% of the surface and that 𝒮_3_ (*t*_2)_ decreases, as 1 / ζ_*n*_ increases. However, the longer (WT) still occupies a significant fraction despite a low initial ratio in the inoculation beads (Supplementary Fig. S18). For instance, when WT is only ∼ 2% of the inoculations (1 / ζ_*n*_ = 47), it still occupies ∼ 6% of the final colonies’ surfaces. Taken together, our results indicate that the longer cell still wins, but that its competitive strength is weakened. This is in accordance with our expectation based on relaxed nematic alignment in 3D.

## DISCUSSION

Using a set of three non-motile *E. coli* substrains, differing only in cell aspect ratios, we revealed that longer bacteria outcompete their shorter counterparts, even when initially outnumbered. By simulating the 2D colony growth where we assumed surface adhesion to dominate on the front, using the agent-based model of growing elongated particles with repulsive interactions, we showed that the advantage originates from local nematic ordering induced by mechanical interactions along the sector boundaries set by the radial expansion. The finding was experimentally confirmed by time-lapses of the takeover process at single-cell resolution: The longer WT cells aligned and formed nematic channels linking the homeland to the expanding front. After this, they aligned perpendicular to the front and eventually surrounded the whole colony, while the shorter AK cells were cut off from nutrients and space. We further confirmed that this advantage is reproduced in 3D colonies, even though the effect was less prominent than in surface environments. We also found (from mono-strain colonies) that WT expands faster over a substrate surface than the shorter mutants (Supplementary Fig. S19+S20), whose slower radial expansion is compensated by an increase in thickness (48–50). As takeover occurs in the monolayer on the expanding front (Fig. S21), the expansion rate is not the determinant of competition, instead it is the strain-strain interaction – enhanced by local alignment. In addition, the WT/AS takeover was significantly slower than for the other combinations (Fig. 3), which cannot be explained from expansion rate differences. Nevertheless, future work could investigate an expansion of the particle model (51), where the expansion of the front occurs on timescales similar to multilayering/verticalization to phase-map the competition between these two effects.

It has been long argued, based on phylogenetic studies, that the morphological development from rod-shaped to round-shaped bacteria (cocci) has happened independently several times and is associated with the loss of *mreB* (52–54). In long-term evolutionary experiments (in liquid medium), an *E. coli* B strain was found to enlarge its volume by shortening and widening (55, 56) in favour of enhanced metabolic efficiency (15). However, our results suggest that on surfaces, the advantageous morphology is the rod shape, enlarging the cell-substrate contact area (50) and colony extension (57) in favour of improved metabolic uptake. In other natural habitats, such as soils and animal tissue (typically associated with spatial constraints and unpredictable supplies of nutrients) the rod-shape can also be advantageous. For example, when communities grow radially in a homogenous environment, as tested in this study.

It is well-understood that morphology shapes colonies (e.g., tortuosity of sector boundaries (Supplementary Fig. S15) (58, 59), swarming efficiency (60), and layering (19)). However, our results highlight an additional, yet so far overlooked, emergent property of bacteria in large communities caused by generic mechanical interactions rather than species-specific chemical cues. The simplicity of this morphological feature suggests a potential to be generalised over a wide range of bacteria species, confirmed by the reproduction of the results using minimal mechanical agent-based model and a generic continuum model. Our findings show how bacterial aspect ratios can influence bacterial advancements in growing colonies. Since biofilm-formation impacts various human activities (61, 62), a better understanding of their mechanical determinants can benefit numerous sectors, including ground remediation, biofouling of industrial installations, and changes in human microbiomes. The mechanical advantage of larger aspect ratios uncovered in this manuscript allows for a generic pathway to control biofilms by modifying the length of various bacteria, which is already achievable with existing technologies (63–65).

## METHODS

### Bacterial strains

We used three subpopulations of the non-motile (66) *Escherichia coli* B strain REL606: the wild-type strain (WT) and the *mreB* mutants (67) REL606*mreB*^*A*53*S*^ (AS) and REL606*mreB*^*A*53*K*^ (AK) (68) (Table 2). All strains are kind gifts from the Kevin Foster Lab. The *mreB* gene in bacteria codes for the filament protein MreB, which is homologous to actin in eukaryotic cells (69). Depletion of the *mreBCD* operon in *E. coli* leads to spherical, enlarged, and eventually lysed cells (70). More specifically, MreB is responsible for maintaining the rod-like cell shape of *E. coli* cells by promoting cell wall growth in regions of negative cell wall curvature. Therefore, its depletion leads to homogeneous cell wall growth (71) by increasing the helical pitch of the filaments, causing shortening and widening of the bacteria’s shape (72). All our strains carried a plasmid that constitutively expressed kanamycin resistance and either GFP, pmaxGFP (pmaxCloning-Vector, Lonza), with excitation/emission of 487/509 nm or RFP, pTurboRFP (pmaxCloning-Vector, Lonza), of 553/574 nm (19).

**Table 2:** *Escherichia coli* strains used in the study.

| Name | Strain | Relevant characteristics | Reference |
| --- | --- | --- | --- |
| WT | REL606 | Ancestral strain. Ara(–), StrepR | (55) |
| AS | REL606*mreB*<sup>A53S</sup> | REL606 with *mreB*<sup>A53S</sup> allele | (68) |
| AK | REL606*mreB*<sup>A53K</sup> | REL606 with *mreB*<sup>A53K</sup> allele | (68) |

### Culture media

Throughout this study, we alternated between the following two media: rich Lysogenic broth (LB) and M63 minimal media (M63+glu) supplemented with 30 µg / ml kanamycin sulphate ( ≥ 95%, K1377, Sigma-Aldrich) to retain fluorescence, unless otherwise stated. For agar plates, we used either 0.5% or 1.5% BD Bacto™ agar (10455513, Fisher Scientific).

#### M63+glu

The 5X M63 salt solution was composed of 15 g/l anhydrous KH_2_PO_4_ (≥98.0%, P9791, Sigma-Aldrich), 35 g/l anhydrous K_2_HPO_4_ (≥99.0%, 60353, Sigma-Aldrich), 10 g/l (NH_4_)_2_SO_4_ (≥99.0%, 09978, Sigma-Aldrich), 2.5 ml 20 mm FeSO_4_ (≥99.5%, 44970, Sigma-Aldrich), 20 mm Na-Citrate (≥99.5%, 71402, Sigma-Aldrich). The M63 minimal media was composed of 20% 5X M63 salt, 1 µg/ml thiamine hydrochloride (≥99%, T4625, Sigma-Aldrich), 2 mm MgSO_4_ (≥99.5%, 63138, Sigma-Aldrich), 2 mg/ml glucose (≥99.5%, G7528, Sigma-Aldrich), dissolved in Millipore water.

#### LB

The medium was composed of 10 g/l Gibco™ Bacto™ tryptone (16279751, Fisher Scientific), 5 g/l Gibco™ Bacto™ yeast extract (16279781, Fisher Scientific) and 5 g/l NaCl (≥99%, S9888, Sigma-Aldrich) dissolved in Millipore water.

### Cell culture

For starting the culture, bacteria from frozen glycerol stocks were streaked on LB or M63+glu agar (1.5%) plates to obtain microcolonies after overnight growth at 37 ^°^C. A single colony from the streak plates was transferred into 2 ml LB and incubated overnight at 37 ^°^C under constant shaking to prevent biofilm formation. The OD_600_ of the overnight cultures was measured after a 10-fold dilution (IMPLEN NanoPhotometer C40) to control the strain proportions in the resulting suspension.

### Imaging

#### Confocal laser scanning microscopy (CLSM)

On a Leica DMI6000 CS SP5 (Leica, Wetzlar, Germany) we used the Ar (488 nm) and HeNe (543 nm) laser lines sequentially to excite GFP (detection: 498 −536 nm) and RFP (detection: 568 −641 nm), respectively. We alternated between the following magnification objectives: 5× air objective (Nplan5×0.12PHO, Leica), 20× air objective (NplaL20x0.40NA, Leica), and 100× oil immersion objective (100×0.17/D/HCX/PL/APO, Leica).

#### Fluorescence microscopy

We used a Nikon ECLIPSE Ti microscope (Nikon, Tokyo, Japan) paired with an Andor Neo camera (Andor, Belfast, UK). GFP and RFP were excited by a Hg lamp using the FITC (FITC Filter Cube Set, Olympus, York, UK) and Texas Red cubes (Texas Red™ Filter Cube Set, Nikon, Tokyo, Japan), respectively. In combination with one of the following objectives: 4× air immersion objective (Nikon, Plan Fluor, 4x/0.13, ∞/1.2 WD 16.5), 20× air immersion objective (Nikon, Plan Fluor, 20x/0.50, ∞/0.17 WD 2.10) and 100× oil immersion objective (Nikon, Plan Apochromat λ, 100x/1.45 Oil, ∞/0.17 WD 0.13).

### Growth rate measurements

Starting from overnight cultures (WT+GFP, WT+RFP, AS+GFP, AS+RFP, AK+GFP, AK+RFP) in M63+glu liquid medium, we diluted 100× in M63+glu to final volumes of 5 ml. The cultures were incubated at 37 ^°^C and the OD_600_ was measured at regular intervals (1.25 h-1.75 h) for 5 − 6 h, while making sure to dilute appropriately to stay within the optimal range of the spectrometer. In parallel, we collected samples for CFU measurements. We diluted the sample to adjust the CFU count to ∼100/plate, and 100 µl was plated on 3 LB agar (1.5%) plates and incubated overnight before CFU were counted.

### Cell aspect ratio

We collected samples (WT+GFP, AS+GFP, AK+GFP) in the stationary phase (i.e. immediately following overnight culture) and the log phase (3 h after dilution) for imaging. Single droplets of bacterial cultures were deposited on cover slides (#1.5) and imaged using fluorescence microscopy and the 100× objective. We measured the length, *l*, and width, *w*, of cells using the line tool in Fiji ImageJ to calculate the aspect ratio.

### Competition in liquid co-cultures

Overnight cultures (WT+GFP, WT+RFP, AS+GFP, AS+RFP, AK+GFP, AK+RFP) were grown in M63+glu liquid medium. The OD_600_ of the overnight cultures was measured and the strains were mixed pairwise (GFP+RFP) in equal proportions (ζ = 1) and diluted 100× in M63+glu. Cultures were incubated at 37 ^°^C and cell counts (50000 counts per sample) were measured right after mixing and again after 8 h of incubation, using a BD FACSJazz Cell Sorter (BD Biosciences, cat. no. 655490). In parallel, we measured the OD_600_ to confirm that the overall growth rates of the co-cultures were the same as for the individual cultures. Data were processed using the Flowkit package (73) with a *logicle* transformation, and clustered with the *hdbscan* algorithm (74). The fraction of double negative events is 6%. After clustering, we extracted the counts of the two main clusters (GFP and RFP) and defined the frequencies of the two populations as the ratio of cell counts of one population (GFP or RFP) over the cell counts of both (GFP+RFP). At *t* = 0, we measured the number ratio of shorter bacteria (mean±SD) to be (29 ± 1)% for AK+WT, (38 ± 0)% for AK+AS, and (44 ± 1)% for AS+WT, when averaging over the colour combinations (*N* = 2). For the AK+WT case, this means that longer cells are ∼ 2.4 more in numbers (compared to ∼ 2.2 from CFU counts).

#### Relative fitness

The relative fitness, *W*_*i j*_, of strain *i* in respect to strain *j* is estimated here as the ratio of the number of doublings of the two competitors (i.e., identical to the ratio of their Malthusian parameters) (55):

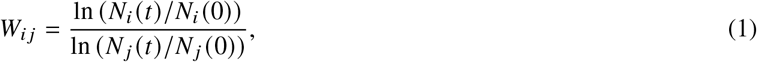

where *N*_*i*_ (*t*) / *N*_*i*_ (0) and *N* _*j*_ (*t*) / *N* _*j*_ (0) are the ratios of number of cells at time *t* in respect to starting point of strain *i* and *j*, respectively. These ratios are obtained from multiplying the ratio of frequencies (i.e., at *t* = 8 h in respect to *t* = 0) with the fold-change in total OD_600_, which in our case was 100.

### Cell density measurements

For each strain (WT+GFP, WT+RFP, AK+GFP, AK+RFP) we plated 100 µl of diluted overnight culture on LB plates with agar (1.5%), which were incubated overnight (37 ^°^C), before CFU were counted. We found similar results for the two colours and measured the (combined) conversion at OD_600_ = 1 to be 1.6 · 10^9^ CFU and 0.7 · 10^9^ CFU, for WT and AK, respectively. Therefore, the cell number ratio between the WT and AK strains in stationary phase (same OD_600_) was measured to be ∼ 2.2.

### 2D competition

#### Sample preparation and imaging

Starting from overnight cultures, we adjusted the ratios (based on OD_600_) of the longer cells (WT or AS) versus the shorter (AS or AK) to ζ ∈ {1, 0.1, 0.01, 0.001} and diluted to a final (and combined) OD_600_ of 0.3. The diluted cell culture was inoculated in 0.5 µl droplets on M63+glu plates with agar (1.5%). After either 20 h or 3 days of incubation (37 ^°^C), we imaged the colonies using the CLSM and the 5× objective. A vertical range of 100 µm was scanned and the resulting 3D images had voxel sizes of (*x, y, z*) = (6.07, 6.07, 10) µm.

#### Pre-processing

In many cases, colonies had outgrown the field of view (3.1 mm × 3.1 mm), so we stitched 4 images together using the Pairwise Stitching plugin in Fiji ImageJ (75). We made a maximum intensity pixel projection of each stack to get a flattened (*x, y*)-image of pixel size (*x, y*) = (6.07, 6.07) µm.

#### Intensity adjustment of colour channels

To boost the contrast between the two channels, we divided channels (GFP and RFP) with each other to amplify the pixel value range of the two channels. Thus, for each pixel, we calculated a new intensity value, *p* (*x, y*)^′^, using a re-scaling value of 100 (8-bit image), such that:

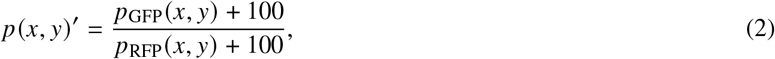

where *p*_GFP_ and *p*_RFP_ are the pixel values from the GFP and RFP channels, respectively. We found this method to be particularly useful in image areas with significant intensity variations (e.g., around the homeland) and for background removal (76).

#### Pixel classification

For further analysis, we wanted all pixels to be classified as either GFP or RFP (never both). Therefore, we used the rescaled pixel intensities, *p* (*x, y*)^′^, distribution of the image and – consistently – found two peaks corresponding to GFP and RFP, and using Otsu’s method, we estimated the threshold value to classify the pixels.

#### Competition strength as a function of the radial vector in 2D

Colonies had clearly detectable regions and we defined the homeland as the well-mixed centre region, which roughly corresponded to where the original inoculation droplet was deposited. The boundary of this region was found by manual fitting of a circle and we identified the centre of this circle as the starting point of the radial vector **r (***t*) ^′^, and the homeland boundary **r (***t* = 0) (after which clear sectors first appear). To account for variations in drop size, we defined the translated radial distance as

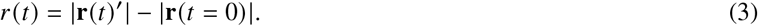

We defined the ensemble-averaged strength of *N* colonies to be the re-scaled occupancy of the GFP-expressing (and longer) bacteria at a given radial length *r* (*t*) as the sum over the following fraction:

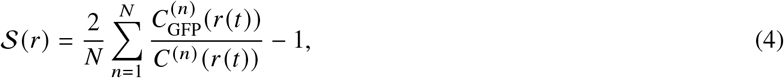

where *C* ^(*n*)^ (*r* (*t*)) is the total number of pixels of the circle of the *n*-th colony at radial distance *r* (*t*) and 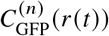 is the number of pixels occupied by GFP-expressing bacteria, such that 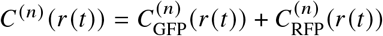. Defined like this, 𝒮 had the value of 1 (-1) when the GFP-expressing (RFP-expressing) cells had taken over the complete perimeter at a specific radial distance. To compare the 𝒮 at a specific time, we defined *t*_1_ = 1.5 days as halfway from the homeland border to the front of a colony after 3 days of incubation.

#### Competition strength as a function of time in 2D

Starting from overnight cultures, we adjusted the ratios (based on OD_600_) of the longer cells (WT) versus the shorter ones (AK) to ζ = 0.001 and diluted to a final (and combined) OD_600_ of 0.3. The diluted cell culture was inoculated in 0.5 µl droplets on M63+glu plates with agar (1.5%). After 11 h of incubation (37 ^°^C), fluorescence images were acquired over a period of 10 h using our fluorescence microscope and the 4× objective. Then, following pre-processing, intensity adjustment, and pixel classification, we measured the time-dependent strength of a colony as the sum over the following fraction:

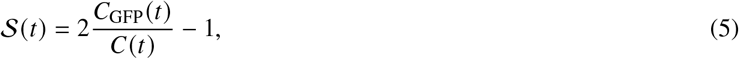

where *C*(*t*) is the total number of pixels of the circle fitted to the periphery at time *t* and *C*_GFP_(*t*) is the number of pixels occupied by GFP-expressing bacteria, such that *C*(*t*) = *C*_GFP_(*t*) + *C*_RFP_(*t*). To calculate *d*𝒮 (*r*)/*dr* we fitted 𝒮 (*t*) for individual colonies and used the *r* (*t*) development at the periphery.

#### Competition in isogenic bacteria

Starting from overnight cultures, we inoculated 1:1 (ζ = 1) two subpopulations (expressing GFP or RFP) of the same strain (WT/WT, AS/AS, or AK/AK) with the same fitness and shape at OD_600_ = 0.3. Plates were incubated for 20 h and CLSM imaged. We found patterns of spatially segregated lineages with a constant number of sectors (Supplementary Fig. S15A); well in accordance with earlier findings (19, 20).

#### Zoom in on 2D competition isogenic boundaries

Plates were incubated (37 ^°^C) for 3 days, before cutting out the colony and the according square of agar and placing it upside down in a glass-bottomed culture well (HBST-5040, WillCo Wells B.V.). The colonies were imaged with sequential z-stacks using CLSM in combination with the 100× objective and the resulting voxel size in (*x, y, z*) = (0.30, 0.30, 0.99) µm.

#### Varying seeding density

We repeated the competition experiment (20 h) for WT/AK but for serial dilutions of seeding density. The co-culture of WT and AK mixed 1:1 and OD_600_ = 0.3 was diluted 10-, 100-, and 1000-fold. We found the mixing of genotypes reduced in the homeland (i.e., patch sizes enlarged), but WT was still able to completely dominate the expansion front (Supplementary Fig. S4). So even though the seeding density regulates patch sizes (77, 78), the longer cell still wins.

#### Rate of range expansion in 2D

Starting from overnight cultures (WT+GFP, AS+GFP, AK+GFP) in LB, we inoculated 0.5 µl droplets of single-strain cultures (OD_600_ = 0.2) on M63+glu agar (1.5%). After 15 h of incubation (37 ^°^C), the plates were placed upside down on our inverted fluorescence microscope and imaged with a frame rate of 1.5 h^−1^ during 9 h using the 4× objective. We mapped the colony areas by (manual) thresholding and quantified the range expansion by calculating the effective colony radius, *r*, from the measured surface area using the Measurement function in Fiji.

#### Cross-sections

Starting from overnight cultures (WT+GFP and AK+RFP) in LB, bacteria were mixed in equal proportions, reaching a final OD_600_ of 0.3. The diluted cell culture was inoculated in 0.5 µl droplets on LB agar plates (1.5%). The plates were then incubated for 20 h (37 ^°^C). Fresh agar (55 ^°^C) was poured into the dishes to fully embed the bacterial colonies prior to sectioning. The agar was poured onto the original agar substrate, between the colonies, to avoid immediate direct contact with the colonies. Dishes were left to solidify at the bench (1 h). Approximately 1 cm^3^ was cut out around the colony, placed on a glass-bottomed dish and cut in half with a scalpel along the diameter of the colony. The halves were then rotated to image the cross-sections. These were imaged at 4× magnification using the fluorescence microscope, as well as 20× and 100× using the CLSM.

#### Monolayer fronts

Starting from overnight cultures (WT+GFP and AK+RFP) in LB, bacteria were mixed 10:1 in favour of AK (ζ = 0.1), before 10-fold dilution, and inoculation on M63+glu agar (1.5%) plates. Following the inoculation, a rectangular piece of agar was cut out, flipped, and transferred upside down into glass-bottomed culture well (HBST-5040, WillCo Wells B.V.) and sealed with Parafilm®. The colonies were incubated (37 ^°^C) for varying incubation times: 7.5 h, 9.5 h, and 10 h. Images were taken of the colony edges with two sequential z-stacks (RFP then GFP) using CLSM and the 100× objective. The z-stacks were taken over vertical ranges of 6 µm to 18 µm, to compensate for slight horizontal tilts of the agar plates. The resulting 3D images had voxel sizes of (*x, y, z*) = (0.30,0.30,0.20) µm.

### Single-cell resolution time-lapses

#### Sample preparation

One colony of each WT and AK was transferred separately into 3 ml LB medium and incubated overnight (37 ^°^C) under constant shaking. The OD_600_ of the overnight cultures were measured and they were mixed in equal proportions (ζ = 1), before 10-fold dilution and inoculation on M63+glu agar (1.5%) plates. Shortly following the inoculation, a rectangular piece of agar was cut out, flipped, and transferred upside down into glass-bottomed culture well (HBST-5040, WillCo Wells B.V.) and sealed with Parafilm®.

#### Single-cell resolution time-lapses

The colonies were imaged in our inverted fluorescence microscope using the 100× objective. Fluorescence images were acquired with a frame rate of 2/h during 12 h. In addition, z-stacks were collected over a total range of 2 µm (0.5 µm spacing) for each time frame to cope with the relative roughness of the agar substrate surface at high magnification.

#### Data analysis

The z-stacks were high-pass filtered and summed into a single image using a custom-made MATLAB procedure, thereby collapsing all the sharpest regions into a single plane. Bacteria were segmented by thresholding each time frame multiple times (twice for AK, thrice for WT) to consider variations in intensity and saturation within a single time frame. The binary images resulting from different thresholds (of the same time frame) were summed to collect as many bacteria as possible, despite non-uniform contrasts, see Movie 6-7 for examples. Bacteria were detected using the Regionprops function in MATLAB (Image Processing Toolbox version 11.7). The function returned the (x,y)-coordinates and the orientation of each bacterium (major axis of the ellipse that has the same second-moments as the bacterium pixels region).

The local nematic order parameter *q* for a bacteria *n*, is calculated by locating the *N*_*n*_ = 8 nearest neighbours *n*_*i*_ of the same strain and then using the definition:

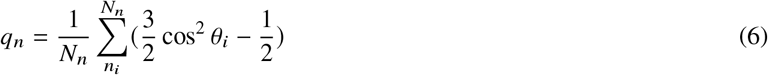

with *θ*_*i*_ the orientation of the neighbouring bacteria orientation. To calculate the probability density function, PDF, all *n* are radially grouped and the front region was defined as the portion for which WT cells form a continuous layer. The intermediate (middle) region corresponds to the AK-rich region of strong segregation behind the front, and the homeland to the disorganized portion further inward. When such regions have not yet formed, the three regions were defined as equidistant radial bands.

### 3D competition

#### Encapsulating bacteria in inoculation beads

We produced four batches of 2.5% agarose inoculation beads containing different ratios of WT+GFP and AK+RFP following the procedure outlined in Ref. (21), excepting the following modifications: Cell densities in the beads were either OD_600_ = 0.65 or OD_600_ = 1.45 (OD_600_ = 4 or OD_600_ = 8.6 in the cell mix) for 1 / ζ_*n*_ ∈ {0.25, 12, 16} and 1 / ζ_*n*_ = 47, respectively. The latter was obtained by up-concentration through centrifugation of the overnight cultures. Varying the densities ensured reasonable numbers of (WT+GFP), despite low ratios (high 1 / ζ_*n*_). We ii) filtered the PBS-bead suspensions through strainers with mesh sizes of 70, 50, and 40 µm and let at least 10 ml additional PBS flow through the strainers to wash off the remaining silicone oil. We collected the beads in the 40 µm strainer (diameters of 40 −50 µm) by turning it upside down and flushing with 6 −9 ml LB. More details can be found in the associated protocol (79).

#### Mono-strain 3D colonies

As a control, we also prepared beads with two colours of the same strain. We followed the exact same procedure but with a different bacteria mixture: Overnight cultures were mixed 1:1 by adjusting the OD_600_ (WT+GFP/WT+RFP, AK+GFP/AK+RFP, AS+GFP/AS+RFP) and diluted in LB to a final (overall) OD_600_ = 0.07 (0.4. in the cell mix).

#### Bead concentration measurements

To measure the strain ratios and number of cells within the inoculation beads, we did serial dilutions (10^−7^ − 10^−5^) of the spare cell mixture from the encapsulation process. We plated 100 µl of the dilution on LB agar (1.5%) plates and incubated overnight. From the number and colour of CFU, we deduced i) the ratio of WT to AK cells (1 / ζ_*n*_) and ii) the overall cell concentration of the mixture. Based on these results, we estimated the number of cells encapsulated within a bead by modelling the beads as perfect spheres with a radius of 45 µm. The strain ratios were estimated from counting 1606, 2900, 3918, and 3472 CFU for 1 / ζ_*n*_ ∈ {0.25, 12, 16, 47}, respectively.

As the small size limited the number of cells per bead and as the strains had different cell volumes (Table 1 in the main text), we defined a strain ratio, ζ_*n*_, as the ratio of CFU of the longer strain versus the shorter (measured before encapsulation in the beads). We found 1 / ζ_*n*_ ∈ {0.25, 12, 16} with OD_600_ = 0.65 and 1 / ζ_*n*_ = 47 with OD_600_ = 1.45; the higher density for the latter ensured WT cells in most beads at this extreme ratio.

#### Embedment of inoculation beads in 0.5% agar

To investigate competition in 3D, we embedded the inoculation beads within a 0.5% agar growth medium. We followed the protocol outlined in (21), excepting the following adaptions: i) Frozen bead stocks were diluted with LB (based on the measured concentration of beads per 1 ml frozen stock) aiming for 10 beads per well. Specifically, we diluted {650,970,130,85 } µl of frozen bead stock in 1 ml LB for 1/ζ_*n*_ ∈ {0.25, 12, 16, 47}, respectively. We counted on average 10.9±3.3 beads per well and ii) we incubated all colonies for 10 h.

#### 3D colony imaging

After incubation, wells were preserved at 4 ^°^C until imaging (< 24 h) using the CLSM and the 20× objective (discarding colonies that had reached the agar-glass or the agar-air interface). For bi-coloured colonies, we recorded two sequential (RFP then GFP) z-stacks with a resulting voxel volume of (*x, y, z*) = 1.51, 1.51, 1.33 µm. For each strain ratio (1 / ζ_*n*_), we collected images from 4-5 different wells (> 40 colonies in total). As a control, we also mixed 1:1 (i.e., ζ_*n*_ = 1) two subpopulations (expressing GFP or RFP) of the same strains and found that surface coverage was similar for both strains (Supplementary Fig. S21). In the case of WT/WT, this is well in accordance with earlier findings (21).

#### Segmentation of 3D images

The 3D colony images were processed, segmented, and filtered for the surface using BiofilmQ (80) and Fiji/Image J (81). Only colonies expressing both strains were segmented, following the exact process described in Ref. (21).

#### Competition strength as a function of time in 3D

We defined the ensemble-averaged strength of GFP-expressing bacteria (WT) on the colony surface as the re-scaled sum over the area fraction:

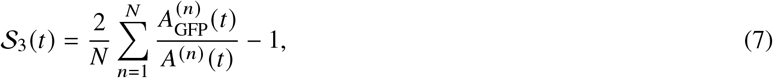

where 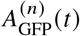 is the area of GFP-expressing bacteria and *A*^(*n*)^ (*t*) is the total surface area of the *n*-th colony at time *t*, such that 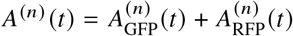. In contrast to the 2D case, 𝒮_3_ is a function of *t* (instead of *r* (*t*)) as it refers to the surface at a given *t*, with varying *r* (*t*). For colonies whose visible surface only expressed one of the two strains, we assigned 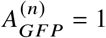 (GFP-mono-coloured) or 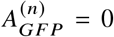 (RFP-mono-coloured). We also calculated the expectation value of the subset of *N* originating from pure RFP-expressing cells from estimated number of cells per bead (CFU/*V*_*bead*_), the total number of colonies (*N*), and the WT/AK ratio (ζ_*n*_); assuming a binomial distribution.

#### Rate of range expansion in 3D

Starting from overnight cultures (WT+GFP, AS+GFP, AK+GFP) in LB, cells were embedded at low concentration ( ∼ 10/well) in M63+glu agar (0.5%) matrix following the protocol described in (11). After 6 h of incubation (37 ^°^C) the mono-clonal colonies were imaged using our fluorescence microscope with a frame rate of 3/2 h during 16 h with a 20× objective. Similar to the 2D case, we quantified the range expansion by manual thresholding and calculated the effective colony radius, *r*, from the (maximum-intensity) z-projected cross-sectional area using the Measurement function in Fiji.

#### Statistics

All mean values are given as mean plus/minus the standard deviation (SD) or the standard error of the mean (SEM), when this is stated and only when data are tested against the null hypothesis that it is normally distributed. The normal distributions of 𝒮 were assessed with Shapiro-Wilk tests. Their distributions were compared between the WT and AK strains with Mann-Whitney U Tests. The significance of the differences in *v* were evaluated with a Students t-test.

### Computational models

#### Particle model

The particle-based model is based on the SPR model presented and described in (8). A single bacteria is described as a rod, *α*, consisting of a rigid chain of a number of Yukawa segments, *n*_*α*_, separated by length, *l* _*α*_, with segments of different rods repelling one another with a Coulomb-like point potential with potential strength, *U*_0_ = 250 (previous work has shown that results are independent on *U*_0_ provided it is large enough (82)), resulting in a total potential, *U*. The Yukawa screening length, λ, is taken as the width of the particle with the aspect ratio of the rod: *L* = *n*_*α*_*l* _*α*_ / λ. Other particle potentials are used in the literature (Bsim 2.0 (83)), but our results are robust independently of this choice.

Bacterial growth is done stochastic. So, for every time step, Δ*t*, we draw a random number **randn** (taken from the absolute value of a normal distribution with mean 0 and standard deviation 1). This random number is multiplied by the current particle length and the growth rate *γ* and then added to the particle length. The choice of distribution and the resulting length polydispersity does not matter. As long as an uncorrelated distribution is chosen, the method in the continuum limit gives the most basic ODE growth solver (Gro (84)):

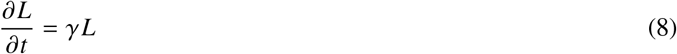

and polydispersity affects the isotropic nematic transition only as a correction. When the rod has reached the division length *l*, the particle is split into two. The timescales given in the manuscript are normalised by the generation time, which is purely set by *γ*.

We use overdamped equations of motion to then update the particle position and orientations (8):

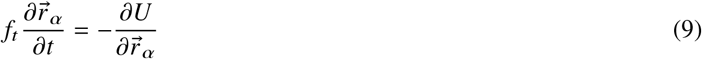

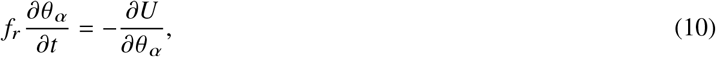

where 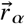 and *θ* _*α*_ are the centre and orientation of rod *α*, respectively and *f*_*r*_, *f*_*t*_ being the drag coefficients. This gives local force propagation even in the non-equilibrium state of an expanding colony, which retains the importance of mechanical interactions that are the highlight of our manuscript. This is in contrast to other bacterial solvers (e.g. Gro (84) and BSIm 2.0 (83)) which use instantaneous global force minimisation to update the bacteria’s positions and orientations.

At time zero (*t* = 0), 40 rods are initialised in a small area with 10 rods having division aspect ratio, *l*_*L*_, and random aspect ratios (uniform distribution) between *l*_*L*_/2 and *l*_*L*_ and 30 rods having division aspect ratio, *l*_*S*_, and random aspect (uniform distribution) ratios between *l*_*S*_/2 and *l*_*S*_, such that ζ = 0.25.

#### Competition strength in the particle model

To determine the competition strength 𝒮 _*f*_ (*t*) at the interface at it evolves over time, we count all particles that exist in the front. We do this by dividing the colony into 10 slices (with centre at the centre of mass of the colony), determine the particle that is furthest from the centre of the colony and then take into account all particles that inhabit up to 5 decay lengths λ into the colony.

The takeover rate, *v*_sim_, in the simulations is calculated from the 𝒮_*f*_ (*t*) curve of 10 different trajectories. The *v*_sim_ is the linear fit from generation time 5.5 to time 7.5.

#### Continuum model

To describe the bacterial colony on a course-grained level, we study the evolution of the nematic order parameter 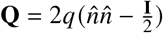, where *q* and 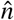 show the magnitude and orientation of the order, respectively, and **I** represents the identity tensor (30).

In line with other dry active nematic studies (85), (43), we use the dynamics of the **Q**-tensor is described by the Beris-Edwards equation

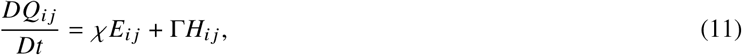

with respect to a symmetric gradients *E*_*i j*_ (shear tensor) and anti-symmetric *ω*_*i j*_ (vorticity tensor) of a velocity field 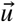. We use the convected co-rotational time derivative *DQ*_*i j*_ / *Dt* = (∂_*t*_ + *u*_*k*_ ∂_*k*)_ *Q*_*i j*_ − *ω*_*ik*_*Q*_*k j*_ + *Q*_*ik*_ω_*k j*_ .

In Eq. 11, *χ* is the aligning parameter (taken as 0.8) and Γ = 0.1 is the rotational diffusivity which along with *H*_*i j*_, the molecular field, describes the relaxation of the **Q**-tensor towards the minimum of a free energy described by:

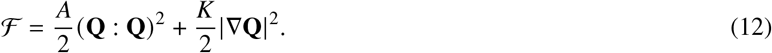

The free energy includes the nematic alignment term (with coefficient *A* = 0.01) and an elastic term *K* which penalises gradients in the **Q** tensor.

Velocity is taken in the overdamped limit

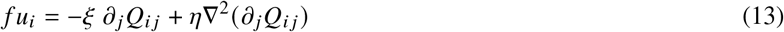

where *ξ* is the activity coefficient related to a force dipoles generated from division events (positive to correspond to the extensile limit), which is balanced by frictional dissipation into the surface *f* and a small viscous dissipation with viscosity *η* = 0.01. The outer edge of the colony is taken by assuming high friction *f* = 400 outside a radius *R*_*con*_ > 30, while lower friction *f* = 4 is used inside the radius *R*_*con*_ < 30 (Supplementary Fig. S10). In addition, the incompressibility constraint is imposed on the total velocity field.

To distinguish the two bacterial concentrations, we introduce a phase-field binary order parameter *𝜙*, which evolves according to a Cahn-Hillard model

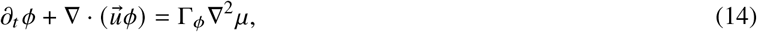

where 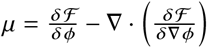 is the chemical potential with Γ_*ϕ*_ = 0.3 a mobility coefficient. We add to the free energy ℱ a single well potential 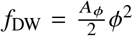 and an interfacial term 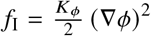 with *A*_*ϕ*_ = 0.002 and *K*_*ϕ*_ = 0.02 such that all bacterial phase-separation comes from activity dependence. We take *𝜙* < 0 as the shorter bacteria and *𝜙* > 0 as the longer bacteria. This is imposed by taking the phenomenological parameters *ξ* and *K* to be dependent on the local *𝜙* as:

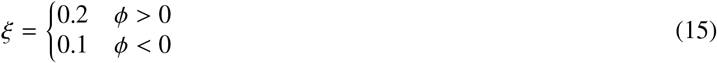

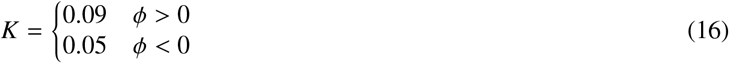

since it is well established that both orientational elasticity *K* and extensile activity due to division *ξ* are higher for higher aspect ratios (38, 39). All equations are solved using a finite-difference method and we initialise our system by taking *𝜙* = 0.01 for radius *R* < 25, between *R* = 25 and *R* = 30 *𝜙* = − 0.01 and *𝜙* = 0 for *R* > 30 in a system of size 80 by 80.

We emphasize that the purpose of the continuum model is to demonstrate how the longer bacteria, through an active anchoring mechanism, can compete at the interface. The model does not capture the initial radial expansion and, as a result, does not produce radially aligned sectors. Instead, the increased activity of the active material enables mixing between species. The phase corresponding to the longer bacteria can advect toward the front, where they align with the interface. This alignment allows them to once again dominate the outer region.

This advective, active mixing mechanism bypasses the need for biomass deposition, which is why the longer bacteria overtake the front in both experiments and particle-based simulations. Recent work on active mixing in particle-based models, driven by reduced friction, shows behaviour consistent with our continuum model, as no radial interfaces are observed (40).

Neglecting the biomass increase is a limitation of the current biphasic continuum model. However, including the biomass effects in the continuum model would require extension to ternary field to allow for an expanding front. Such an extension could capture the impacts of radial expansion together with active anchoring effects between the two expanding species in the bulk.

#### Radial induced alignment due to growth

The alignment mechanism at the interface between two species can be understood in terms of active stresses, expressed as ∇(*ξ Q*). At an interface characterized by a unit normal vector *m*, the dominant contribution to the force density arises from 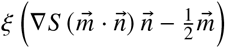, where ∇*S* represents gradients in the nematic order. For simplicity, we neglect gradients in *ξ*; their inclusion would only amplify this effect. Importantly, this interfacial force vanishes only when the director field *n* is either tangential or parallel to the interface. Otherwise, the resulting force generates a flow that reorients *n*, tangentially in extensile clusters or normally in contractile ones (24).

We emphasize that at the front of the colony, this active force will force the longer bacteria to become parallel aligned with this front, which allows the longer bacteria to spread along the interface.

For single-species colonies, we observe no global radial alignment, consistent with earlier observations (Supplementary Fig. S15 and Fig. S16) (43, 44). In radial expanding colonies without perturbations, the outward flow generated by growth alone does not produce sufficient strain or vorticity to drive large-scale radial alignment. Instead, we find that local patches of nematic order emerge spontaneously, but these domains remain uncorrelated with each other and with the overall growth direction. Previous work (44) demonstrated that deliberately breaking radial flow symmetry (i.e. through voids) can facilitate global radial alignment. Our findings suggest that gradients in the local nematic order itself are enough to break symmetry, reinforcing nematic alignment within the ordered regions and allowing for localized self-organization without requiring external perturbations.

## ACKNOWLEDGEMENTS

The authors thank Dr. Michael Lisby for access to the FACS instrument and for generously sharing his expertise. This work was supported by the Independent Research Fund Denmark grant no. DFF0165-00032B (LJ) and grant no. DFF0165-00103B (LJ). The Novo Nordisk Foundation grant no. NNF18SA0035142 (AD), NERD grant no. NNF21OC0068687 (AD), grant no. NNF21OC0068775 (NM), Emerging Investigator grant no. NNF23OC0085012 (KT), and Synergy grant no. NNF23OC0086712 (LJ). The Villum Fonden Grant no. 29476 (AD), and the European Union via the ERC-Starting Grant PhysCoMeT (AD) and the Horizon 2020 research and innovation programme Marie Sklodowska-Curie grant no. 101029079-SIMMS (KT).

## AUTHOR CONTRIBUTION STATEMENT

NM and LJ conceived the original idea. NvdB and TTN conducted the experimental research including analysis. Specifically, NvdB did the surface-attached colony investigations and TTN the 3D investigations. AD conceived the cell-based and continuum models. KT did the cell-based and continuum modelling. MC and AGV co-supervised the project and conducted substantial control experiments. NM, AD, and LJ supervised the project.

## COMPETING INTEREST STATEMENT

The authors declare no competing interests.

## Notes

### Competing Interest Statement

The authors have declared no competing interest.

### Summary of Updates

small typos and corrections Materials and Methods included to the main manuscript

