## Supplementary figures and images for "Emergent collective alignment gives competitive advantage to longer cells during range expansion"

### Movie 1

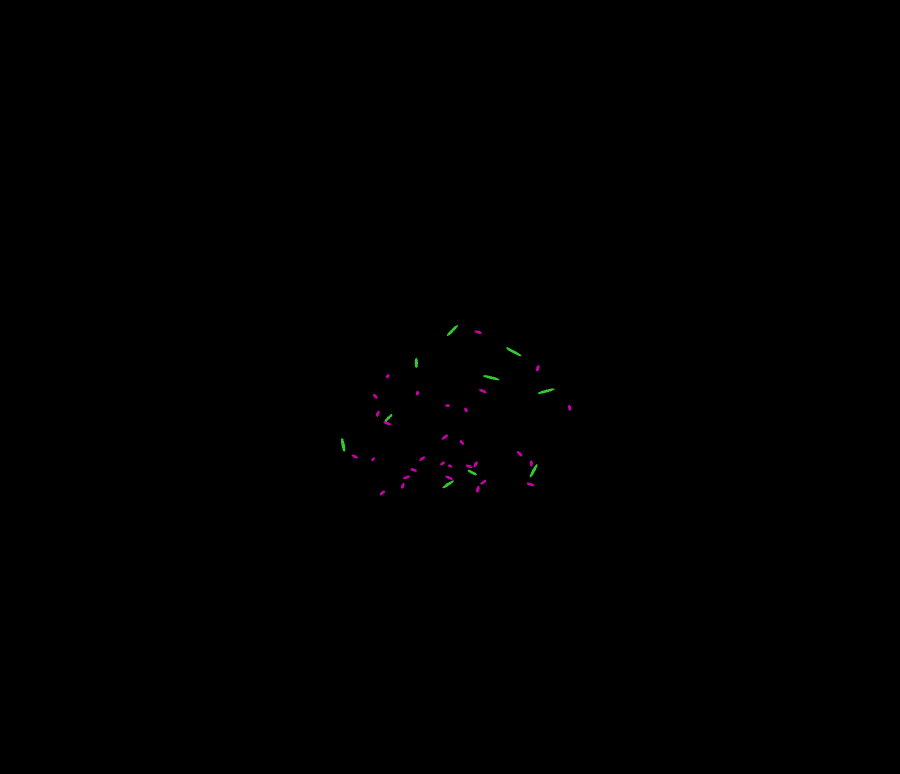

### Movie 2

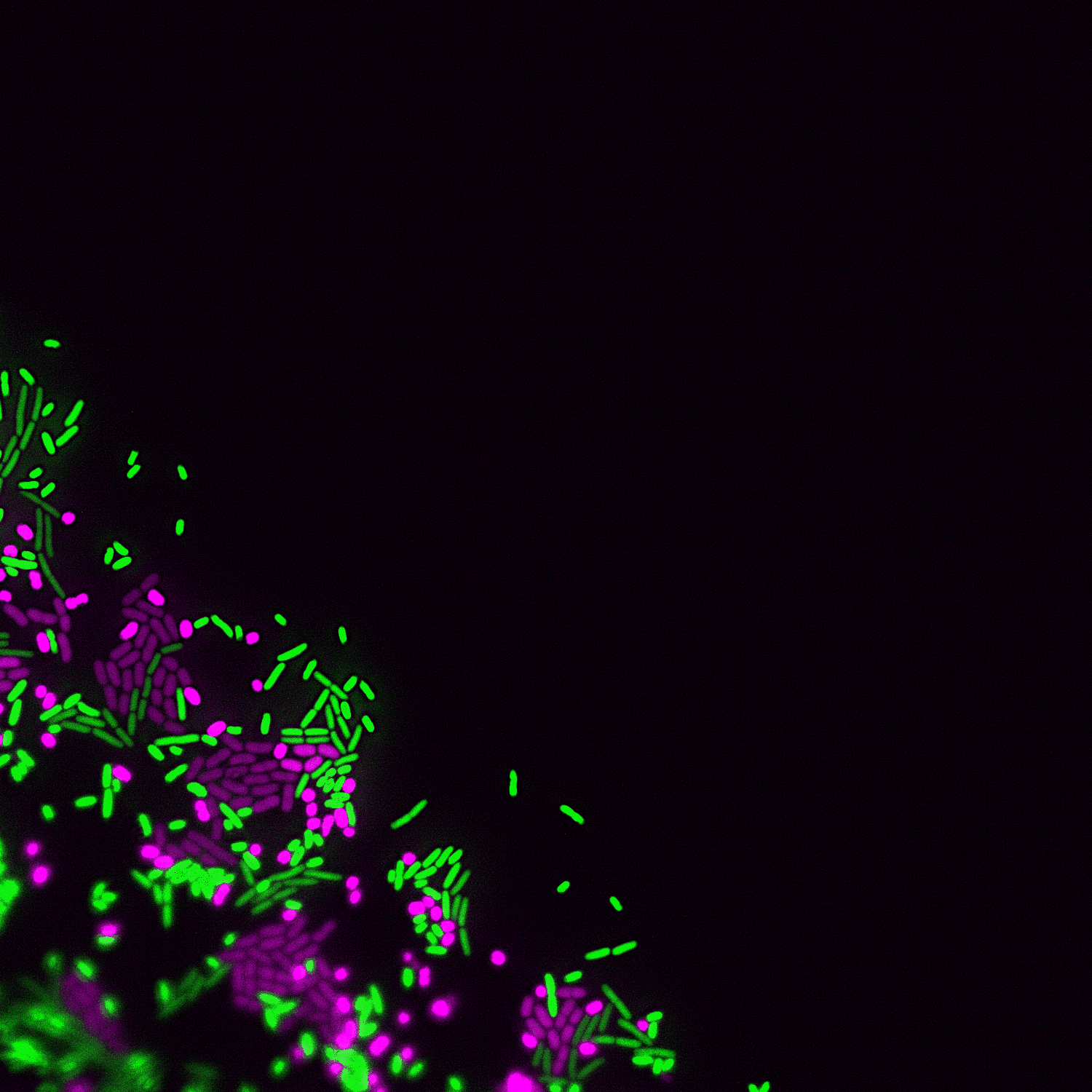

### Movie 3

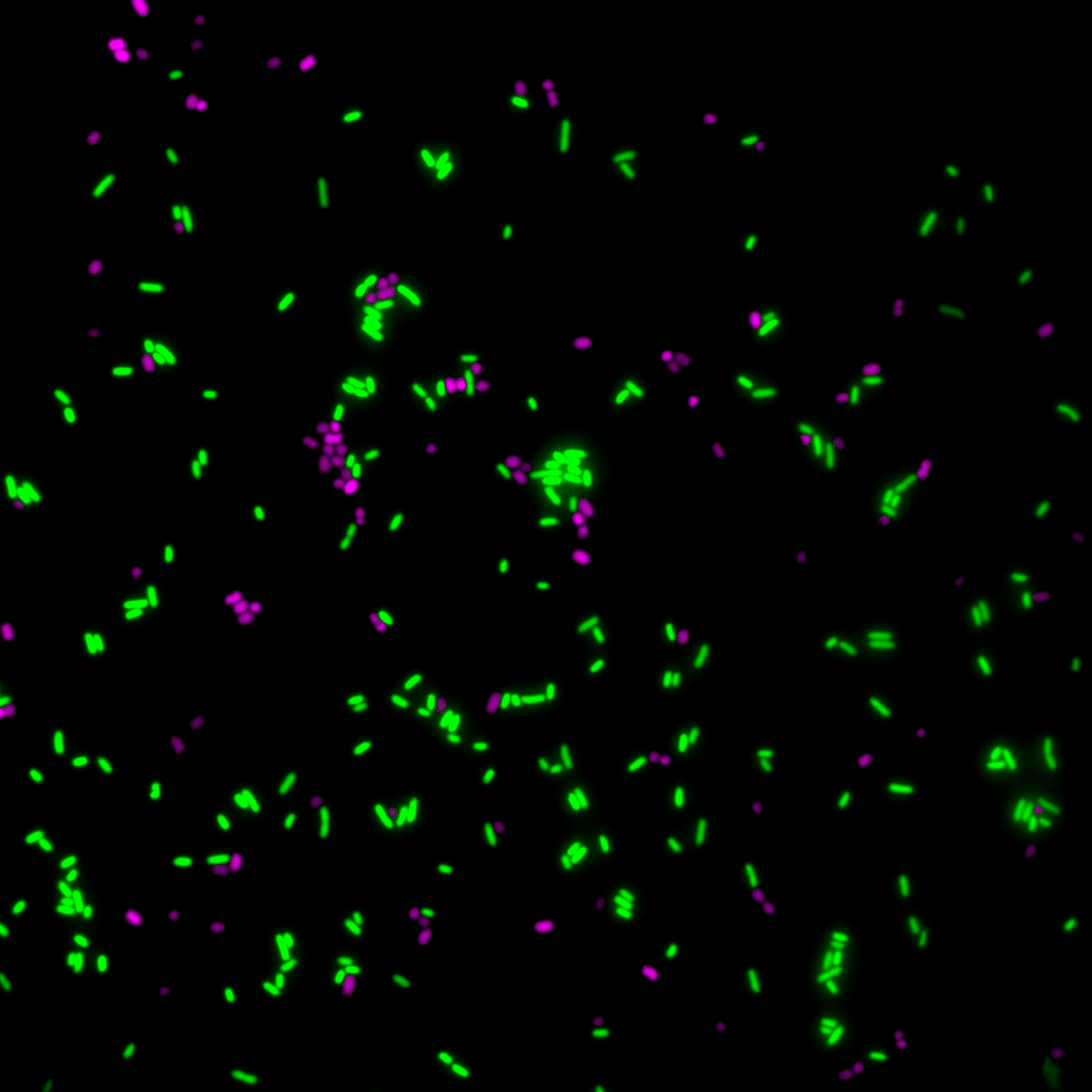

### Movie 4

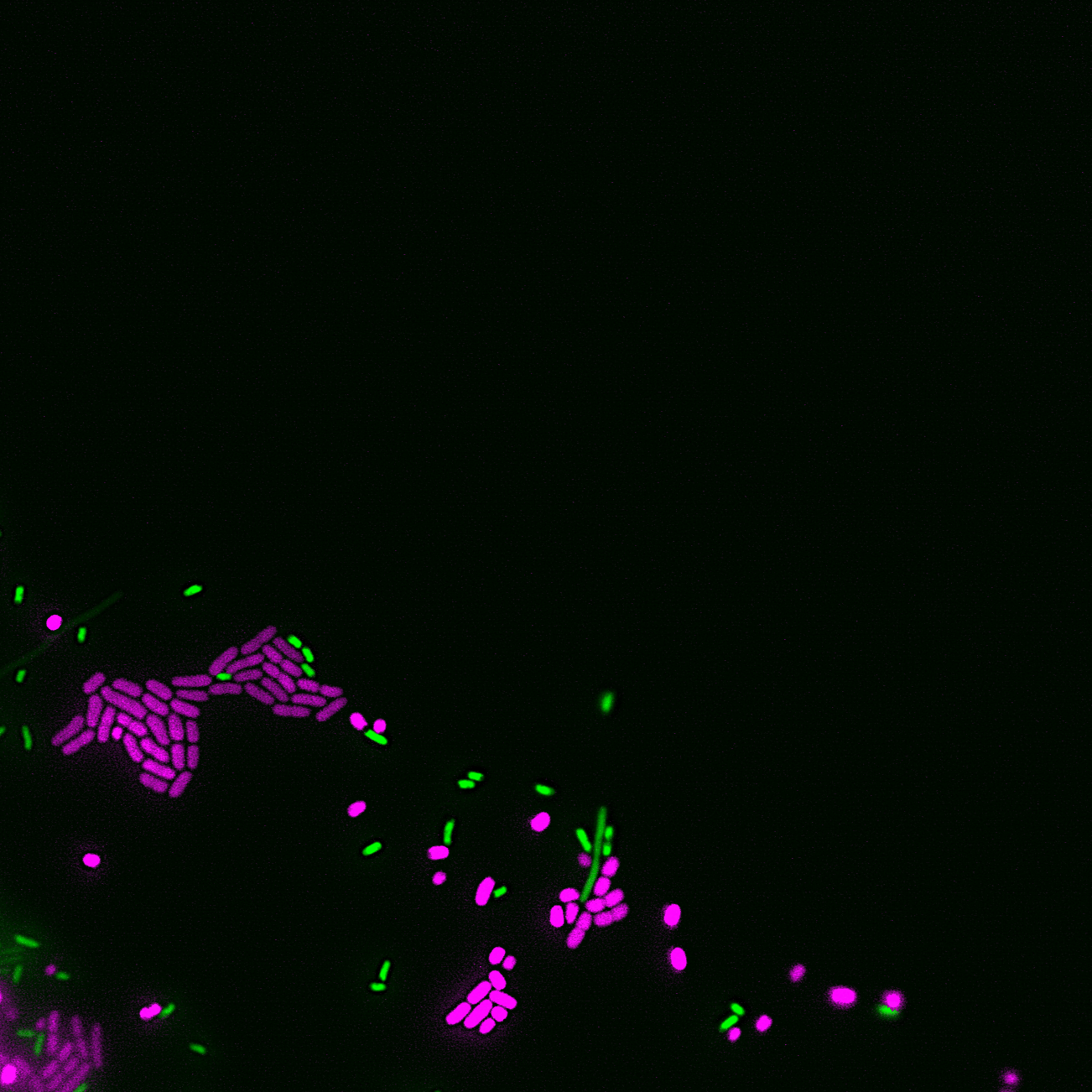

### Movie 5

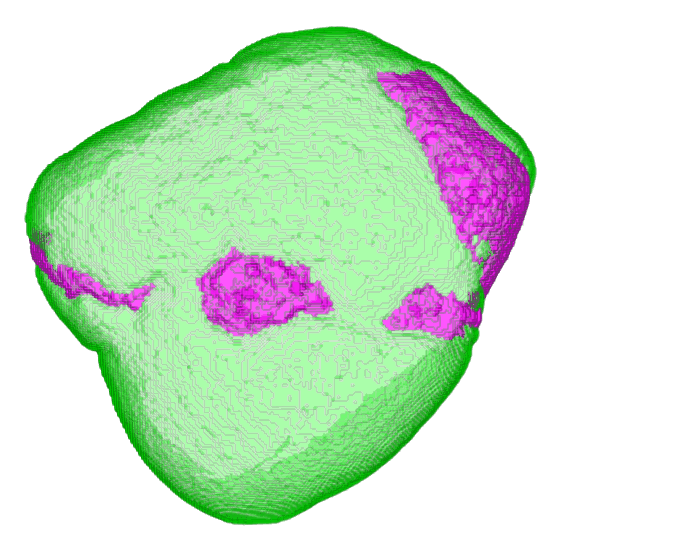

### Movie 6

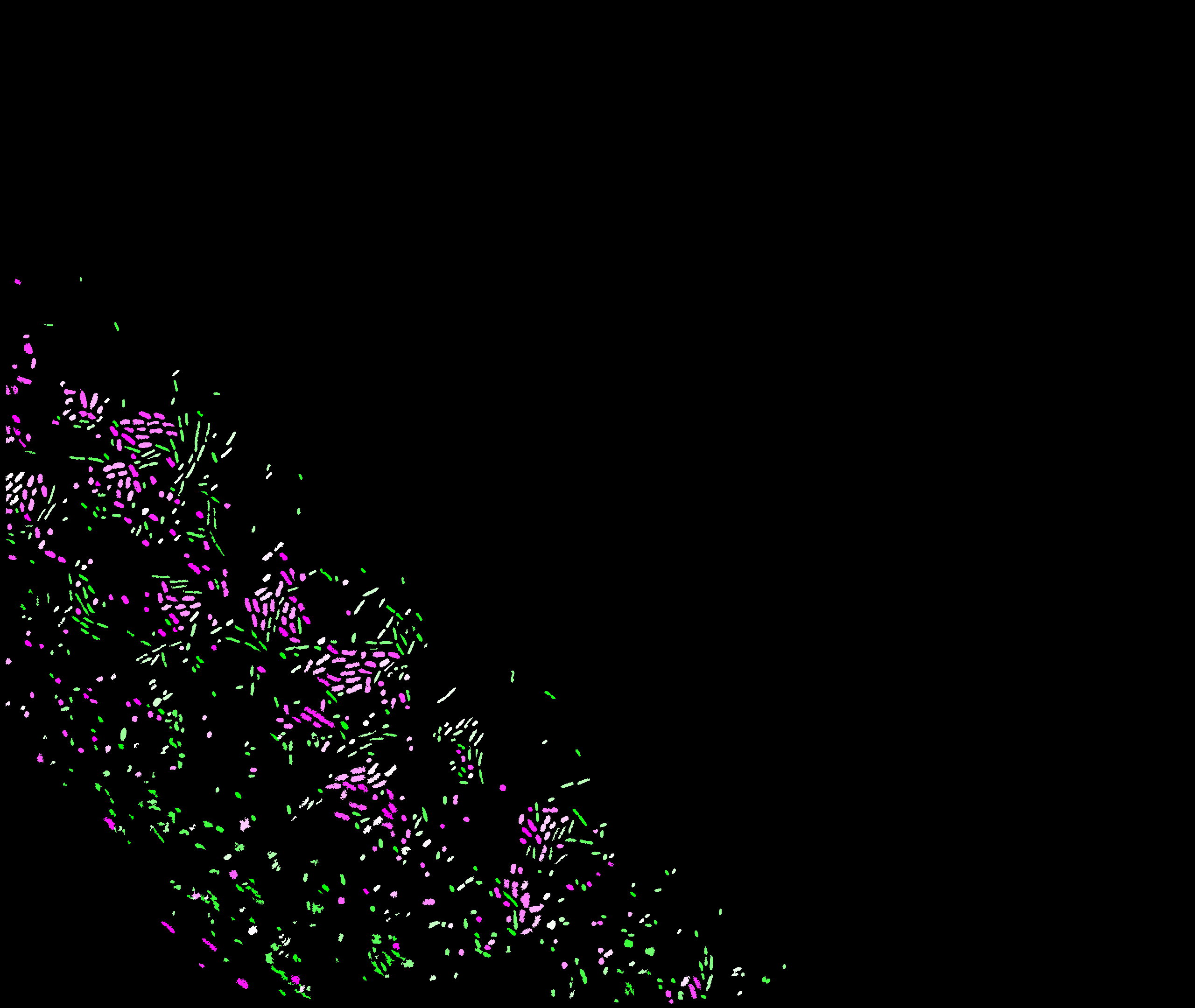

### Movie 7

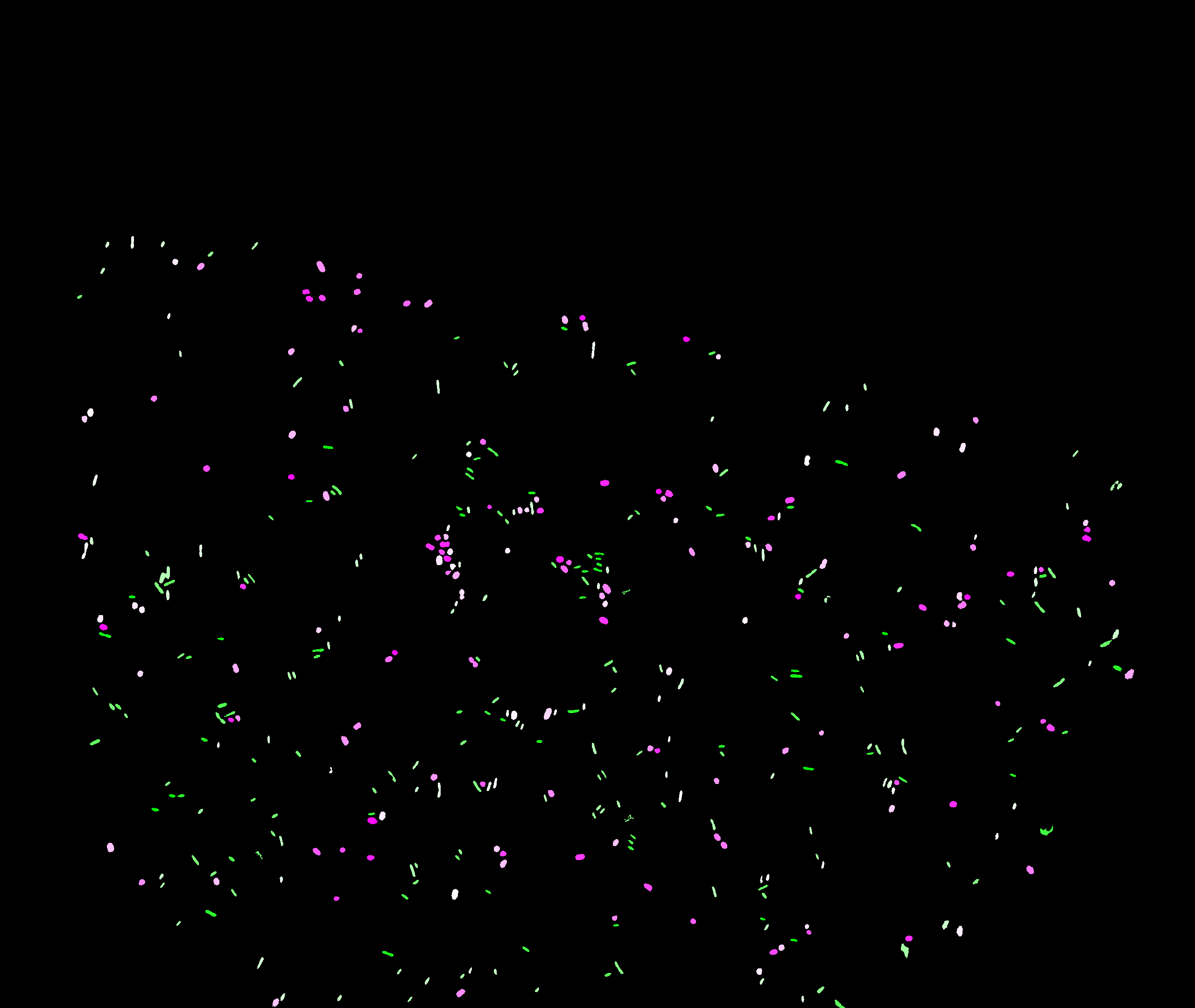
