## Supplementary Information for "Emergent collective alignment gives competitive advantage to longer cells during range expansion"

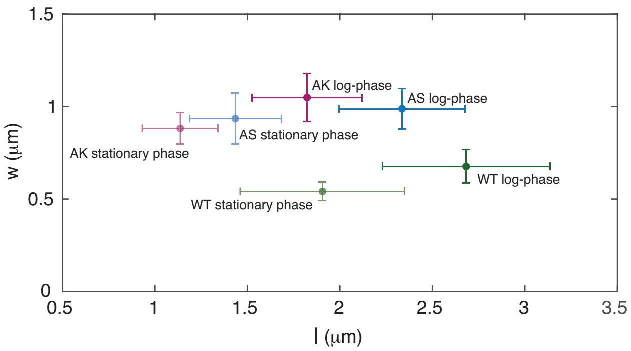


**Figure S1**. Aspect ratios for all strains used in this study. Width, $w$, versus length, $l$, of WT in both stationary phase ($N = 32$) and log-phase ($N = 39$), AS in both stationary phase ($N = 47$) and log-phase ($N = 38$), and AK in both stationary phase ($N = 25$) and log-phase ($N = 33$). Here stationary phase is measured directly in the overnight culture (0 h) and the log-phase 3 h after dilution (Fig. S2). The error bars are ±SD.

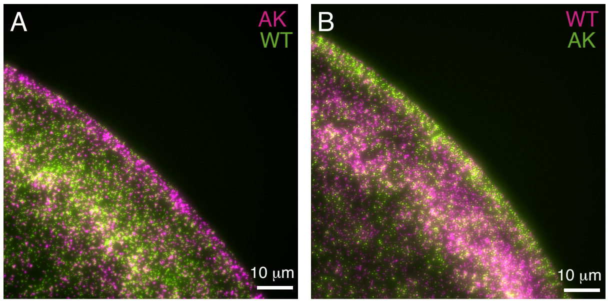


**Figure S2** Fluorescence images (pseudo-coloured) of example "coffee rings" of dried inoculations. Starting from overnight cultures, we adjusted the ratios (based on $OD_{600}$) of the longer cells (WT) versus the shorter (AK) to $\zeta = 1$ and diluted to a final (and combined) $OD_{600}$ of 0.3. The diluted bacterial culture was inoculated in 0.5 μl droplets (at room temperature) on M63+glu plates with agar (1.5%) and imaged instantly at 20× magnification. The scale bars correspond to 10 μm. **A-B**: Concentration of bacteria on the rim of the inoculation droplet: WT (green) and AK (magenta) (A) and swapped colours WT (magenta) and AK (green) (B).

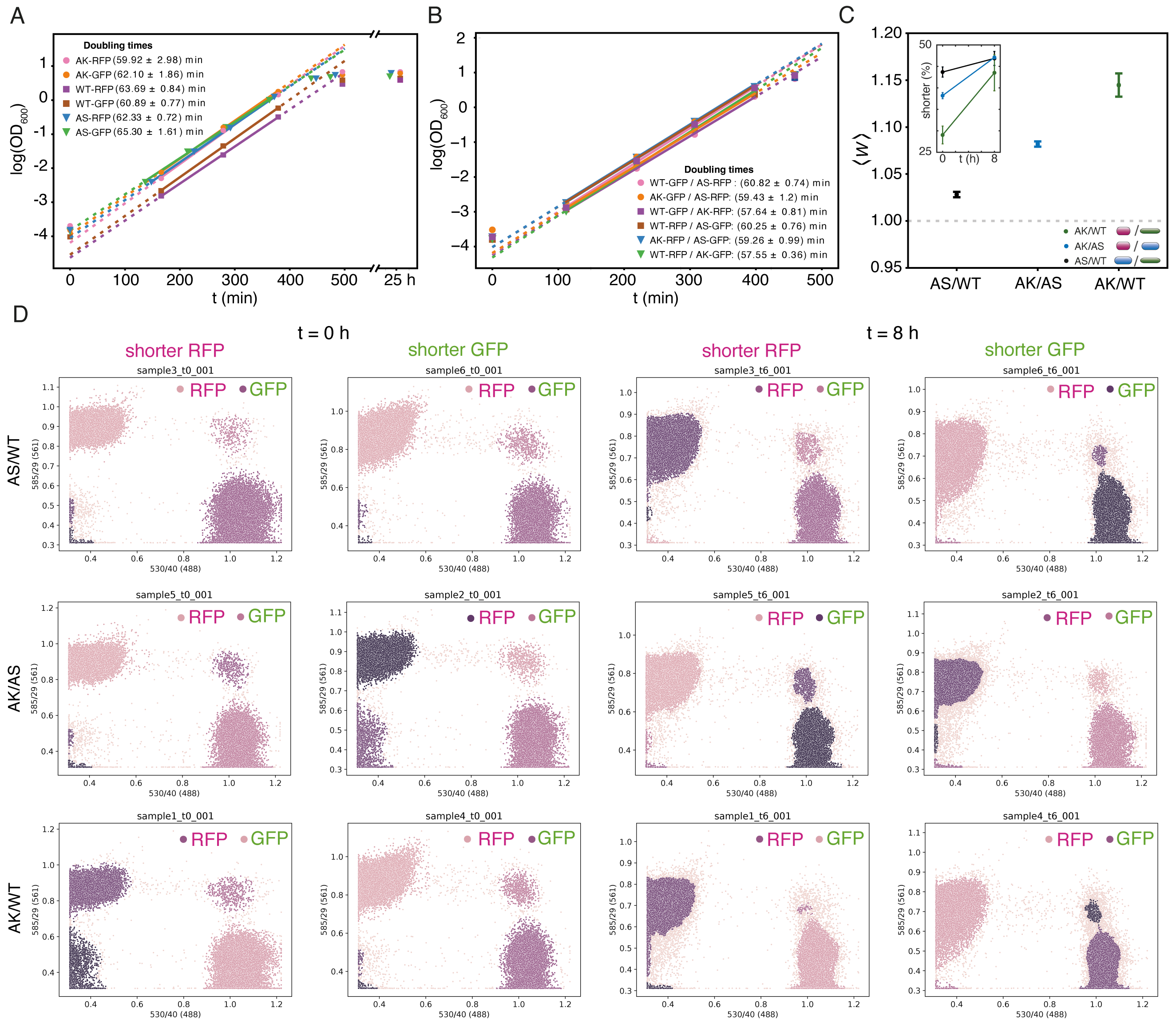


**Figure S3**. **A:** Growth rates for all strains used in this study. The optical density, $OD_{600}$, versus time, $t$, on a semi-logarithmic scale. The solid lines are linear fits to the data and the legends state the corresponding doubling times ±SD. **B:** Growth rates for all strain combinations used in this study (same cultures as in (C)). The optical density, $OD_{600}$, versus time, $t$, on a semi-logarithmic scale. The solid lines are linear fits to the data and the legends state the corresponding averaged doubling times ±SD. **C:** Pairwise competitions ($\zeta = 1$) between all strain combinations in liquid medium (M63+glu) under constant shaking (D). The average fitness of the shorter strain relative to the longer, $\langle W \rangle$, is the ratio of doublings after 8 h averaged over the two independent cultures ($N = 2$) with inverted fluorescence colouring. Error bars are ±SD and the horizontal dashed line corresponds to equal fitness ($\langle W \rangle = 1$). The trend is the same as earlier reported (1). Note that initially the longer cells were more abundant in numbers with a factor ∼ 2.4, ∼ 1.6, and ∼ 1.3 for AK+WT, AK+AS, and AS+WT, respectively. The inset shows the fractions of shorter cells (shorter), which is either AK or AS (see legends). The fractions are averaged over independent cultures ($N = 2$) with inverted fluorescence colouring and error bars are ±SD. **D:** FACS data clustering to obtain the ratio of doublings (C), using fluorescence marker gating and bulk sorting, as detailed in Methods. Source data are provided as a Source Data file (2).

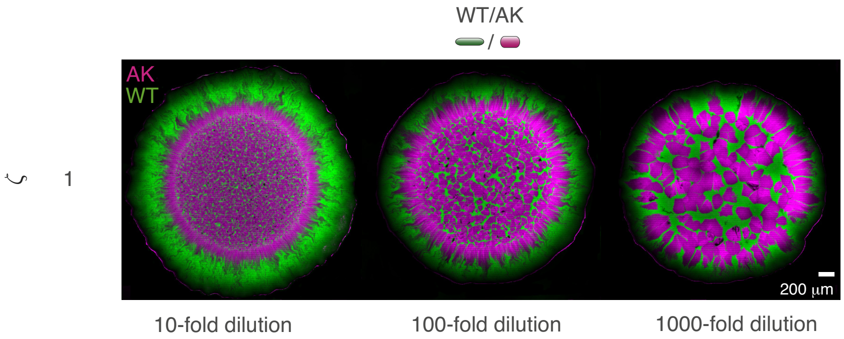


**Figure S4**. Seeding density regulates patch sizes but still the longer cell wins. Competition experiment (20 h) for serial dilutions of the optical density, $OD_{600}$, in a co-culture of WT and AK mixed 1:1 ($\zeta = 1$) and diluted **A:** $OD_{600}/10$, **B:** $OD_{600}/100$, and **C:** $OD_{600}/1000$. For the latter, narrow channels of WT (green) are detectable between the patches of AK (magenta).

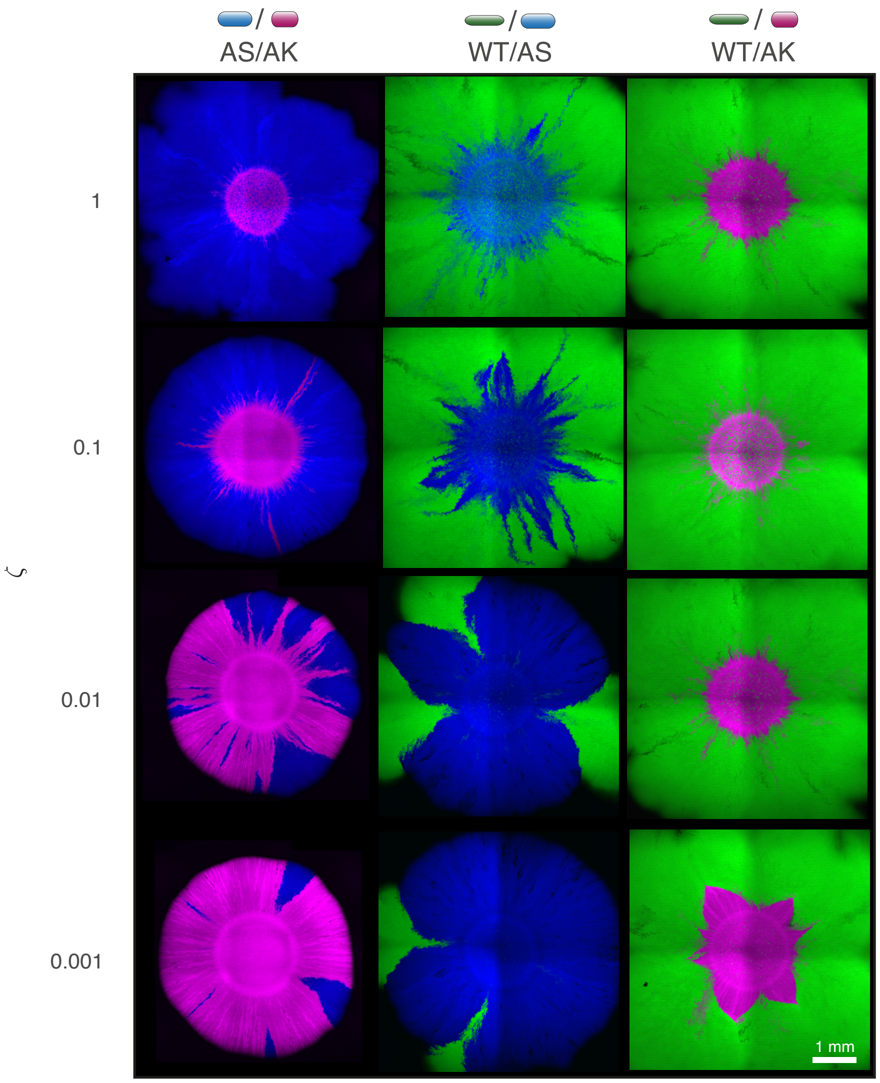


**Figure S5**. 3 days competition experiments. Fluorescence images (pseudo-coloured) example colonies (maximum intensity) z-projection from all strain combinations (AS/AK, WT/AS, WT/AK) imaged by CLSM. The scale bar corresponds to 1 mm. For WT/AK, with $\zeta = 0.001$, we only found 14 AK sectors surviving for 1.5 days (halfway from homeland to front) in 5 colonies, which closed off (i.e., sector was lost) rapidly hereafter. We note that even when the longer bacteria did not overtake the shorter within the explored time window, the sectors with longer bacteria tended to spread azimuthally with increasing $r$, indicating that they might eventually find their way to the expanding front.

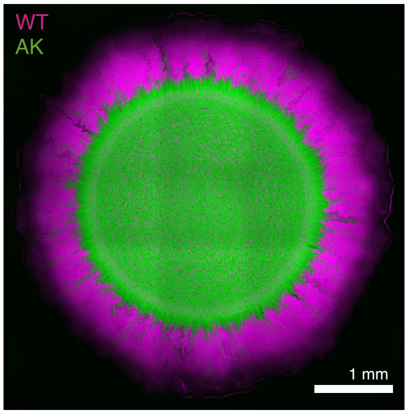


**Figure S6**. 20 h competition experiment with inverse colours. Example colonies (maximum intensity) z-projection from 1:1 ($\zeta$ = 1) WT/AK combination imaged by CLSM. The scale bar corresponds to 1 mm.

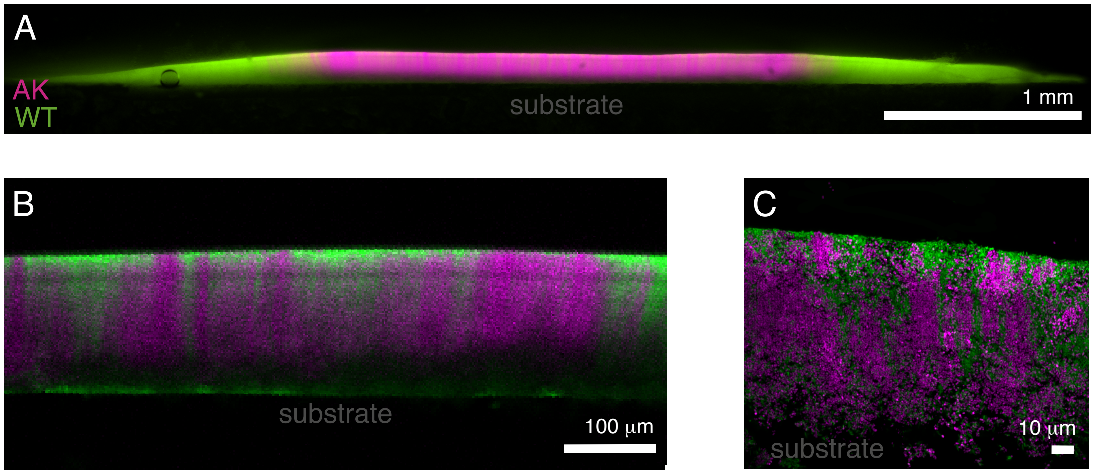


**Figure S7**. Cross-sections of 20 h competition experiments of WT (green) and AK (magenta) colonies ($\zeta$ = 1). **A:** Example colony imaged with fluorescence microscopy and the 4× air objective. The scale bar corresponds to 1 mm. In the outer bands (i.e., outside of the homeland) the number of sectors are constant. **B-C:** Single layer of (pseudo-coloured) CLSM images from the centre of the colony (homeland) using the 20× air objective and a scale bar of 100 µm (B) and the 100× oil immersion objective and a scale bar of 10 µm (C). To contrast these cross-sections of the homeland (B-C), see Fig. S20 for high-resolution images of the monolayer front.

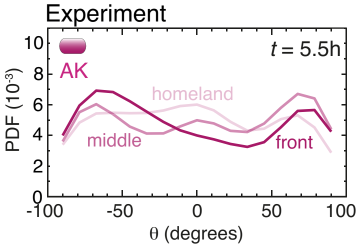


**Figure S8**. The probability distribution function, PDF, of the angle, $\theta$, to the expansion front at three equidistant regions of the radius, $r$: where the inoculation droplet originally was deposited (homeland, $N$ = 239), closest to the outer rim (front, $N$ = 324), and the region in between (middle, $N$ = 687). Data is the AK bacteria at $t$ =5.5 h and from Fig. 2C. Note how AK bacteria are less radially (tangentially) aligned in the middle (front) region than WT (Fig2D).

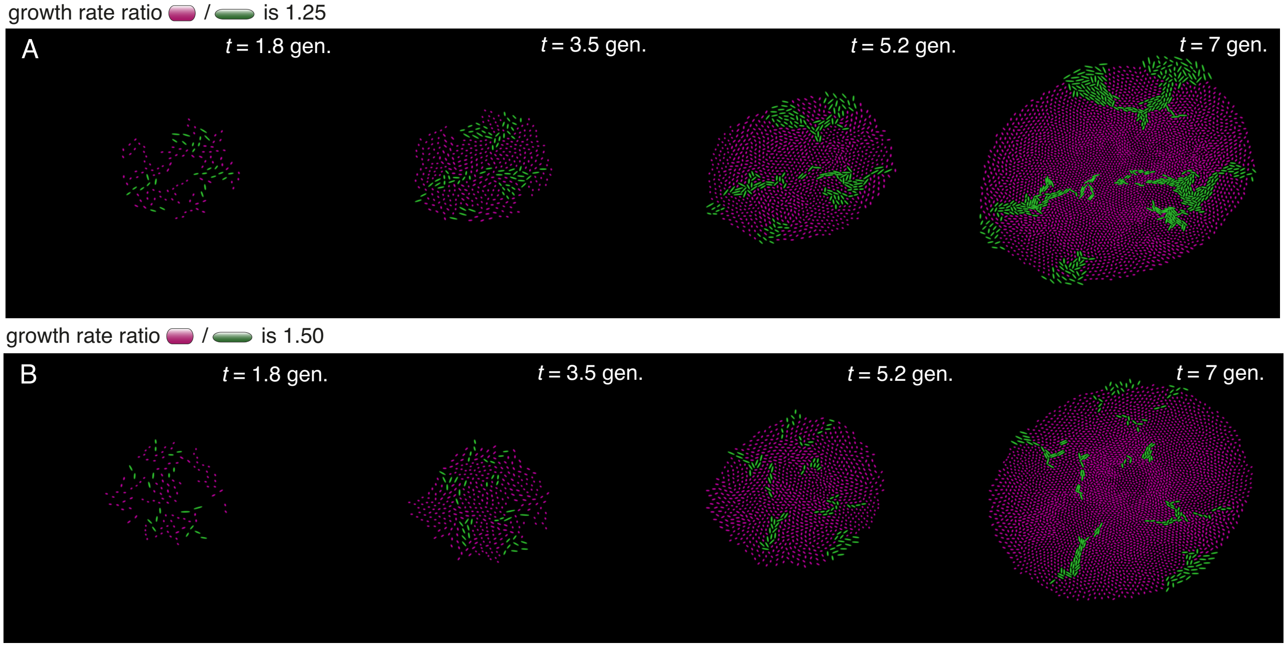


**Figure S9**. Representative snapshots of the simulation where the growth rate, $\gamma$, of the shorter bacteria (magenta) is faster than the shorter (green). **A-B:** Representative snapshots from the simulation where shorter particles divide 1.25× (A) or 1.5× (B) faster than the longer bacteria and time is normalised with the generation time (gen) of the shorter. We remark that numbers start to dominate over mechanical interactions, such that even if the longer bacteria do overtake regions at the interface, the larger bacteria do not dominate the front, as the distance between long-dominated regions expand exponentially as smaller bacteria double (A). For even larger differences in growth rates (B), the number of shorter bacteria grows even more rapidly and the enlarged pressure in the smaller bacteria's sectors results in the longer bacteria being swapped away.

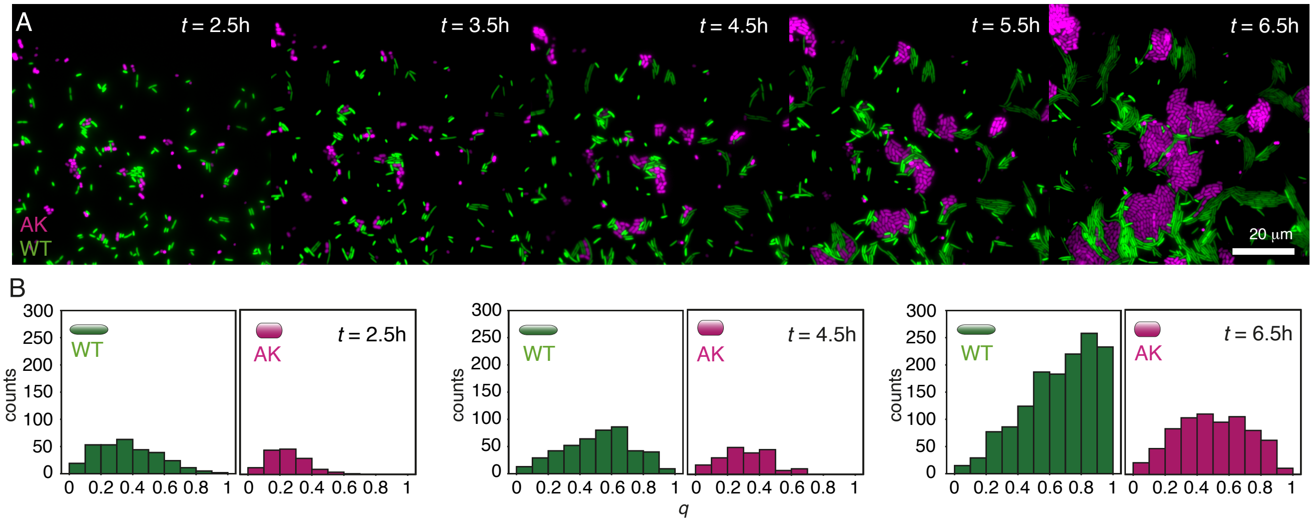


**Figure S10**. Long bacteria align more within clusters. **A:** Single cell (pseudo-coloured) time-lapses of competition between the two-colour wild-type (WT) and the *mreB* mutant (AK) with $\zeta = 1$ at time points, $t$. Representative region of the larger images from which the data plotted in B was computed. The scale bar is 20 μm. **B:** The nematic order parameter, $q$, for WT and AK individually corresponding to every second time frame in (A) $t \in \{2.5, 4.5, 6.5\}$h.

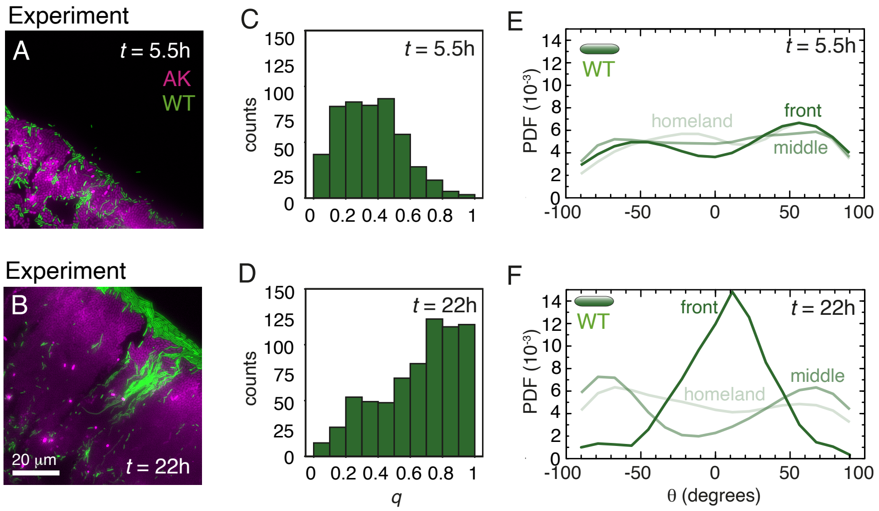


**Figure S11**. **A-B:** Snapshots from (pseudo-coloured) fluorescence time-lapses of competition between WT (green) and the *mreB* mutant AK (magenta) at time $t$ =5.5 h (A) and $t$ =22 h (B) after inoculation. The scale bar corresponds to 20 µm. **C-D**: The nematic order parameter, $q$, of WT corresponding to $t$ =5.5 h (C) and $t$ =22 h (D). **E**: The probability distribution function, PDF, of the angle, $\theta$, to the expansion front at three equidistant regions of the radius $r$: where the inoculation droplet originally was deposited (homeland, $N$ = 51), closest to the outer rim (front, $N$ = 46), and the region in between (middle, $N$ = 391). Data are from $t$ =5.5 h. **F**: As in (E) but at $t$ =22 h and (homeland, $N$ = 202), (middle, $N$ = 254), and (front, $N$ = 242). Note how WT bacteria at $t$ =5.5 h and lower density (E) are less radially (tangentially) aligned in the middle (front) region than later, at $t$ =22 h (F).

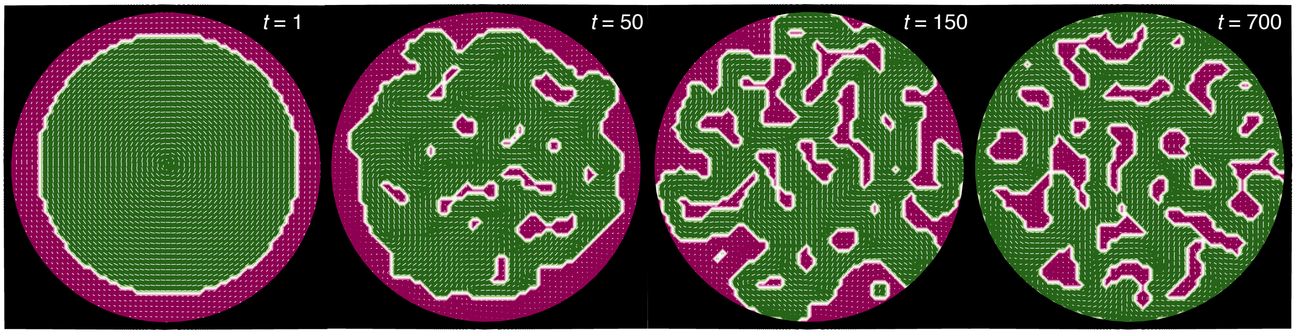


**Figure S12**. Time series of continuum dynamics where the phase representing short bacteria (magenta) starts at the edge while with time, the phase representing long bacteria (green) overtakes the short bacteria by forming pathways to the colony front.

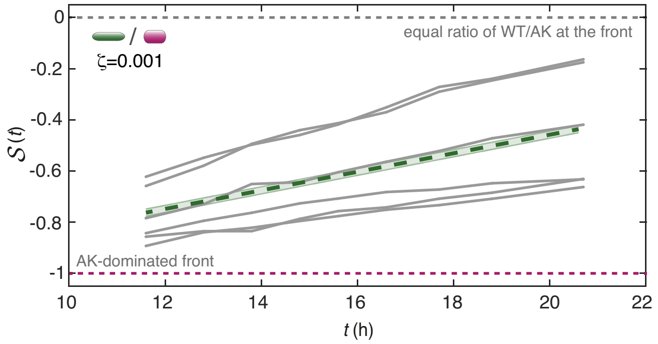


**Figure S13**. Time dependence of the takeover in the case of WT/AK at $\zeta = 0.001$. Competition evaluated by the strength at the periphery, $\mathcal{S}(t)$, over time, $t$, for individual colonies (grey) and the ensemble average of the linear fits (green). The shaded regions correspond to ±SEM ($N = 6$). Note how $\mathcal{S}(t)/dt$ (slope of green line) corresponds to the takeover rate, $v = (1.2 \pm 0.2) \cdot 10^{-3}$ µm$^{-1}$, when the change of colony radius $r$ over $t$ is used to convert the unit. This is well in agreement with the results in Fig. 3B.

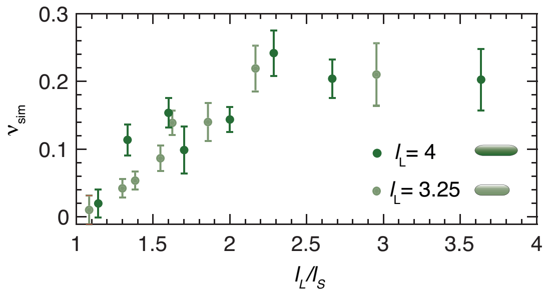


**Figure S14**. The takeover rate, $v_{\mathrm{sim}}$, (slope between $t = 30$ gen. to $t = 45$ gen.) for varying length ratios, $l_L/l_S$ bacteria ratios, where long bacteria still have $l/w = 4$.

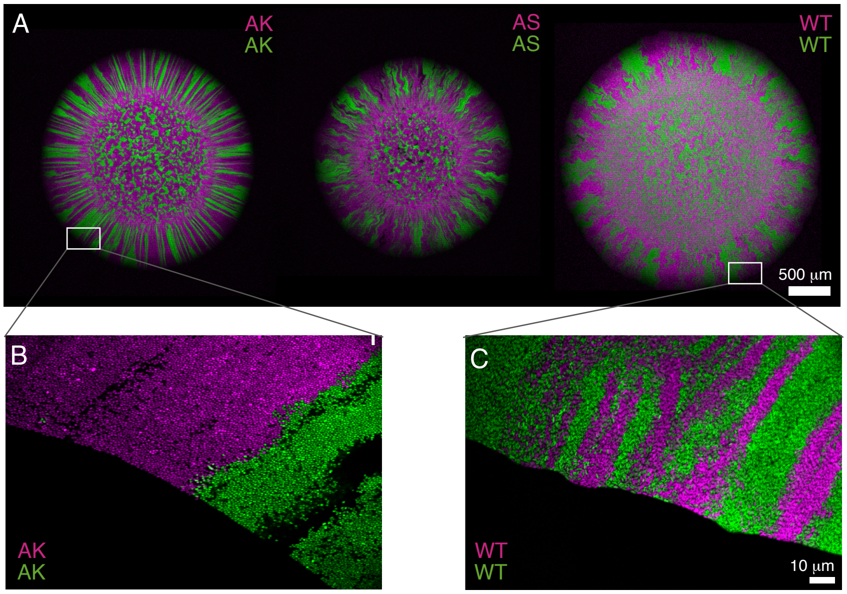


**Figure S15**. Segregation patterns vary with genotype. **A:** The (maximum intensity) z-projections of example colonies (WT/WT, AS/AS, and AK/AK) imaged by CLSM after incubation (20 h). The scale bar corresponds to 500 $\mu$m. In the outer bands (i.e., outside of the homeland) the number of sectors are constant. Despite the general features, qualitative differences among the strains are clear: WT's sector boundaries are coarse (highly diffusive) and the mutants have straighter (less diffusive) boundaries. **B-C:** Single layer (pseudo-coloured) images of competition between the two-colours at the front of the colony: The *mreB* mutant (AK/AK) in (B) and wild-type (WT/WT) in (C). The scale bar is 10 $\mu$m.

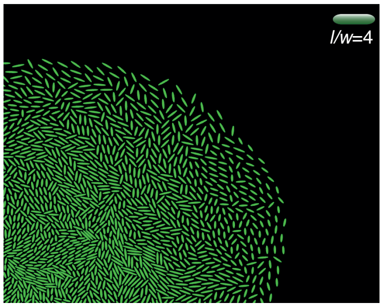


**Figure S16** Representative snapshots of a mono-strain simulation with longer bacteria (aspect ration $l/w = 4$) after 7.1 generations. We remark that there is less nematic alignment than in the simulations with $l_L/l_S = 4$ (Fig. 2A), only small patches of local nematic order where observed with no global radial symmetry.

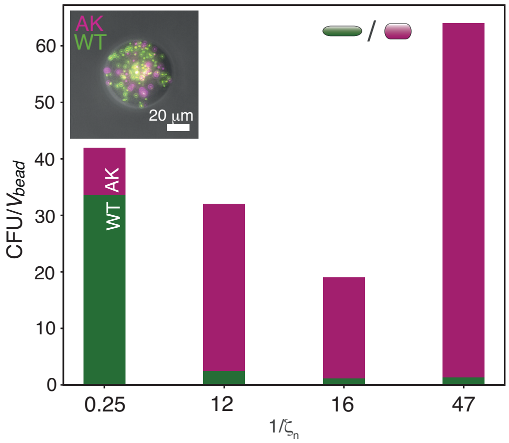


**Figure S17**. Cell concentrations in the various bead batches. The ratio of colony-forming units, CFU, versus the average bead volume, $V_{bead}$, for each strain ratio, $1/\zeta_n$. The colours indicate the fraction of WT (green) and AK (magenta). Inset: Merged image of the bright-field (grey), WT (green), and AK (magenta) of a bead with $1/\zeta_n = 0.25$. The scale bar corresponds to 20 µm.

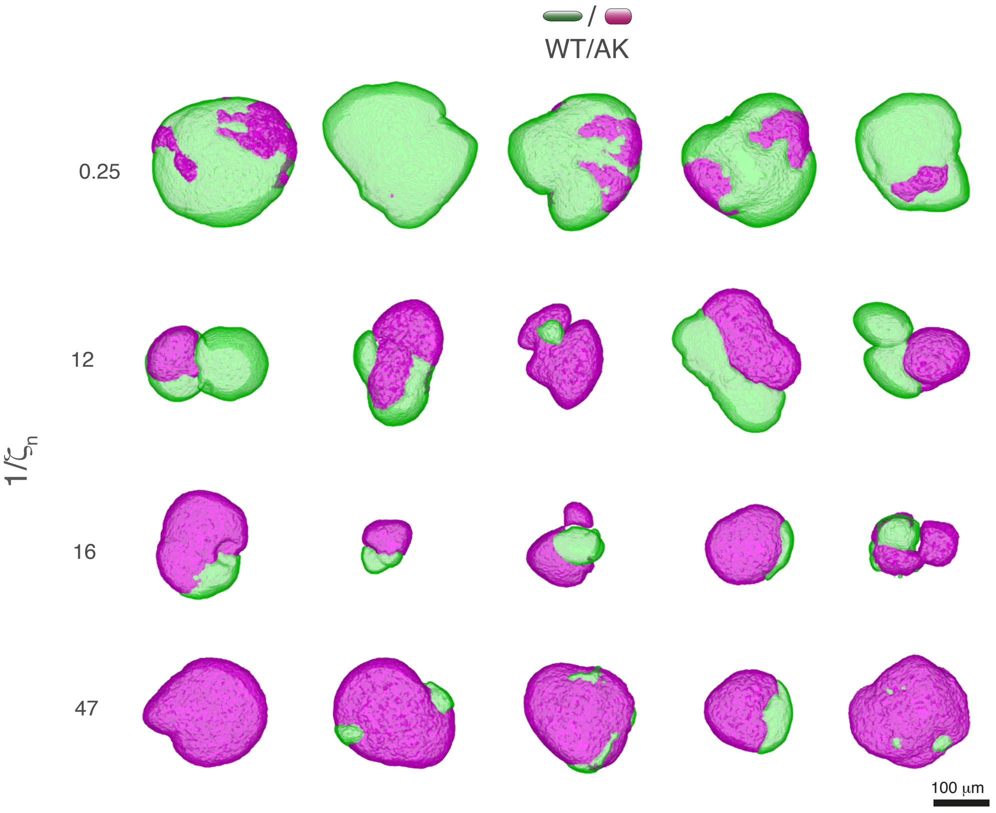


**Figure S18**. 10 h competition experiments. Example segmented masks of surfaces of two-coloured WT/AK colonies from all strain ratios imaged by CLSM. The scale bar corresponds to 100 µm.

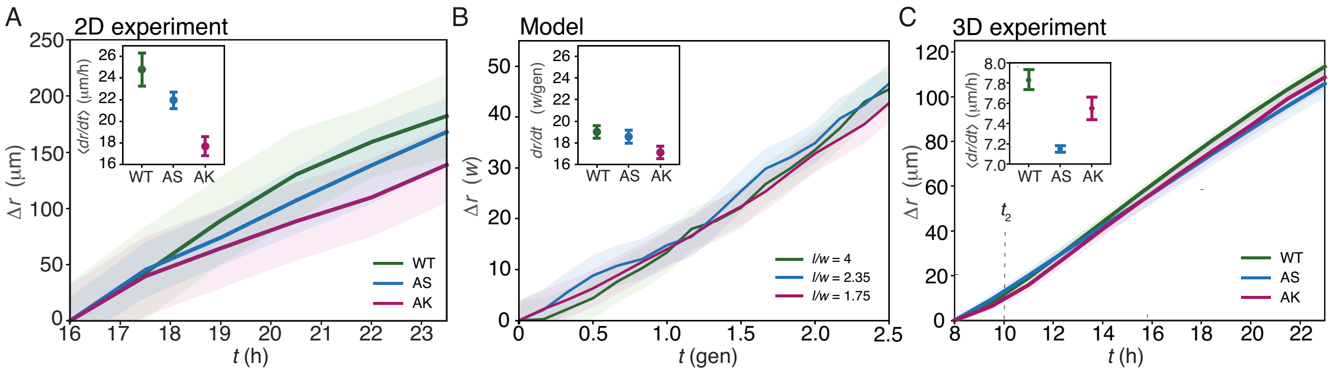


**Figure S19**. Long bacteria spread faster over substrates. **A:** Ensemble-averaged radial expansion, $\Delta r = \langle r(t) - r(16\,\mathrm{h})\rangle$, over time, $t$, of mono-strain colonies of WT ($N = 4$), AS ($N = 6$), and AK ($N = 5$) on agar (1.5%) substrate (M63+glu). The shaded regions correspond to ±SEM. Inset: The ensemble-averaged expansion rates, $\langle dr/dt\rangle$, are linear fits of the data and the error bars are ±CI. **B:** Ensemble-averaged radial expansion, $\Delta r = \langle r(t) - r(0)\rangle$, of simulation data from mono-strain inoculations of cells with aspect ratios: $l/w \in \{4, 2.35, 1.75\}$. $N = 10$ colonies for each $l/w$. The shaded regions correspond to ±SEM. Inset: The ensemble-averaged expansion rates, $\langle dr/dt\rangle$, in number of cell widths, $w$, per generations (gen) are linear fits of the data and the error bars are ±CI. To compare (A) and (B), 1 gen is approximately 1 h and $w$ slightly less than 1 μm. **C:** 3D radial growth of mono-clonal single strain colonies. Ensemble-averaged radial expansion, $\Delta r = \langle r(t) - r(8\,\mathrm{h})\rangle$, versus time, $t$, for WT ($N = 5$), AS ($N = 6$), and AK ($N = 6$). The shaded regions correspond to ±SEM and the vertical punctuated line to $t_2 = 10\,\mathrm{h}$. The inset is the ensemble-averaged slopes of the linear fits, $\langle dr/dt\rangle$, and the error bars are ±CI. Note that this is in accordance with earlier findings (3, 4).

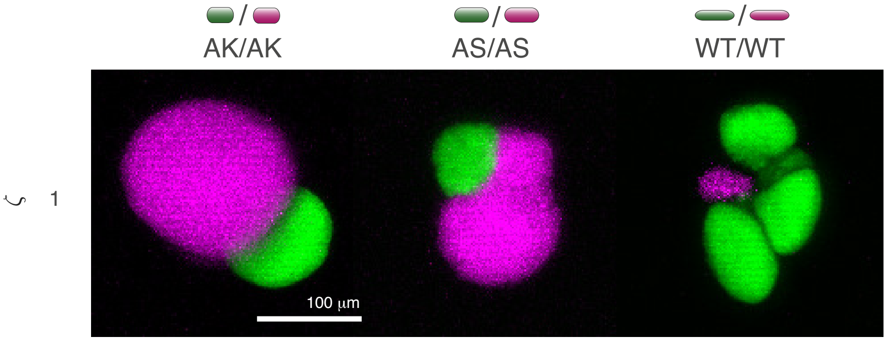


**Figure S20**. 10 h competition experiments. Example colonies (maximum intensity) z-projection from two-coloured strains of AK+AK, AS+AS, WT+WT ($\zeta = 1$) imaged by CLSM. The scale bar corresponds to 100 μm.

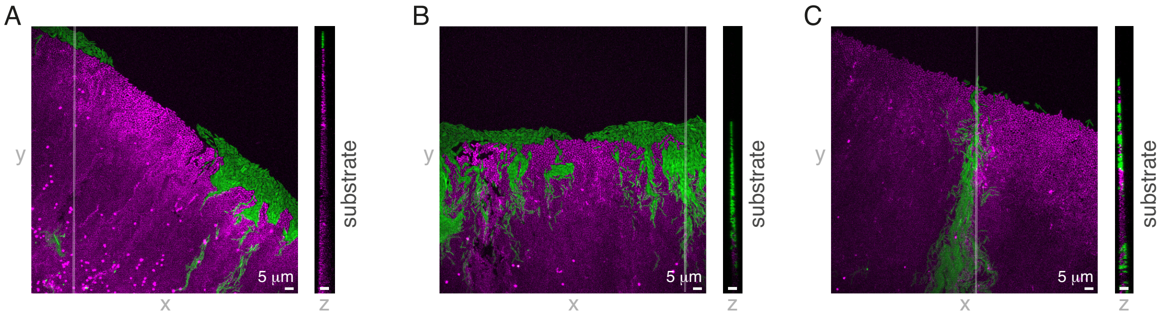


**Figure S21**. Long bacteria win in a monolayer front even when they are disadvantaged. A-C: Maximum-intensity projected (pseudo-coloured) confocal scanning laser microscopy images of takeover events from competition ($\zeta = 0.1$) between WT (green) and the mreB mutant AK (magenta) at time $t \in \{7.5, 9.5, 10\}$ h after inoculation. The opaque lines in the $(x, y)$-view (right) designate the $x$ of the $(y, z)$-view (left) and the scale bars are 5 µm. Note that the monolayer spans over several generations of bacteria at the front.
